# Single residues within the YTHDF intrinsically disordered region control m^6^A-dependent mRNA decay

**DOI:** 10.64898/2026.09.18.752746

**Authors:** Benedetta Ricci, Danika Sommer, Daymieri Narvaez, Ruqi Zhang, Jacob Geri, Sara Zaccara

## Abstract

Intrinsically disordered regions (IDRs) mediate protein interactions, condensate partitioning, and regulatory control in RNA-binding proteins, yet the residue-level logic underlying IDR function has remained difficult to define using conventional fragment- and truncation-based approaches. Here, we apply cytosine and adenine base-editor screens to map residue-level regulation within the intrinsically disordered region of YTHDF2, a cytoplasmic reader that couples m^6^A to mRNA decay. By tiling base edits across YTHDF2 in a YTHDF1/YTHDF3-null background, we identify IDR residues that regulate YTHDF2-dependent cellular fitness. Interestingly, we identified residues that limit YTHDF2 decay activity. Their mutation disrupts endogenous protein-protein interactions and generates hyperactive YTHDF2 variants that promote m^6^A-dependent mRNA decay and alter recruitment to cytoplasmic RNA granules, without affecting intrinsic m^6^A binding or protein stability. These findings define the residue-level regulatory logic of the YTHDF2 IDR and establish base-editor screening as a strategy to uncover functional mechanisms encoded within IDR regions.

## INTRODUCTION

More than 40% of the coding transcriptome undergoes the addition of one or multiple methyl modifications of adenosines, also known as m^6^A ^1–5^. From early studies in the late 1970s to recent transcriptome-wide analyses, mRNA degradation has emerged as a major mechanism through which m^6^A controls RNA fate ^6,7^.

This degradation process is mediated by the cytoplasmic m^6^A readers YTHDF1, YTHDF2, and YTHDF3 (YTHDF1/2/3; DFs), which act redundantly to promote decay of m^6^A-modified RNAs ^8–11^. Although the reader proteins and downstream decay outcomes of this pathway have been extensively characterized, the mechanisms that tune the efficiency of YTHDF-dependent degradation remain poorly understood.

YTHDF proteins share a conserved domain organization composed of a C-terminal YTH domain and a largely intrinsically disordered N-terminal intrinsically disordered region (IDR) ^12^. The YTH domain provides the biochemical basis for m^6^A recognition, and structural studies have defined how conserved aromatic cage residues selectively bind methylated RNA ^12–15^. In contrast, the molecular logic by which the IDR regulates YTHDF activity remains limited.

The IDR has been proposed as the principal regulatory interface that tunes YTHDF function, coordinating protein-protein interactions and recruitment of effector pathways that shape m^6^A-dependent RNA fate^11,16,17^. Consistent with this model, several studies have linked the YTHDF IDR to core RNA-decay machinery, including the CCR4-NOT scaffold CNOT1 ^8^, UPF1 ^18^, and HRSP12 ^19^. However, YTHDF IDR interactions are not restricted to decay factors. YTHDF proteins also associate with proteins involved in RNA granule biology and mRNA stabilization^11^, including the stress-granule scaffold G3BP1 ^20^ and FMRP^21^.

Despite the growing list of IDR-associated proteins, the specific residues or motifs within the IDR that mediate these interactions have not been systematically mapped, leaving uncertainty on how the IDR discriminates between decay-promoting and mRNA stabilization interactors at the single-residue level. To identify the IDR sites that regulate these interactions, the dominant strategy has been to generate fragment-level truncations and then quantify how these IDR perturbations reshape YTHDF interactions and downstream decay activity. Truncation-based analyses have thus mapped N-terminal regions required for association with core decay and surveillance factors mainly with YTHDF2, including the CCR4-NOT scaffold CNOT1 (aa 169-200) ^8^ and UPF1 (aa 101-168) ^18^, and the RNAse P endoribonuclease RNA decay pathway (e.g., HRSP12) (aa 1-100)^19^.

However, there are key limitations within this approach. Firstly, many of these IDR-regulatory fragments are identified using bacterially expressed truncation constructs and in vitro binding assays, making it difficult to determine whether the same region governs YTHDF interactions and function in living cells ^8^. Secondly, when cellular studies have been performed, these have often relied on overexpressed tagged deletion constructs ^18,19^, which do not preserve endogenous protein levels or intact IDR architecture. This is particularly important for IDR, where large truncations can disrupt the spacing and multivalent binding of short linear motifs, alter local charge patterning, and remove flanking sequence context required for motif function, thereby obscuring the interpretation of which specific interactions are regulated ^22,23^. In addition, because YTHDF paralogs act redundantly in mammalian cells, functional effects of perturbing one paralog can be masked by compensation from the others. Thus, although previous studies established that YTHDF IDRs are important regulatory platforms, they have not defined how regulatory information is encoded within the intact IDR at residue-level resolution.

To overcome these limitations, here, we used cytosine and adenine base-editor screens to introduce coding substitutions across full-length YTHDF2 in a YTHDF1/YTHDF3-null background. This sensitized genetic background removes paralog compensation and makes cellular fitness dependent on the remaining YTHDF2 protein, allowing individual YTHDF2 edits to be linked to quantitative cellular phenotypes. Because base editors preserve full-length protein architecture while introducing precise nucleotide substitutions, this strategy enabled residue-level interrogation of IDR function in living cells ^24–26^. By tiling coding edits across YTHDF2, we identified IDR residues whose mutation alters YTHDF2-dependent cellular fitness. We then further validated Tyr68 and Tyr153. Strikingly, mutation of either single residue was sufficient to convert YTHDF2 into a hyperactive degradation mutant. Mechanistically, they disrupt YTHDF2 association with regulatory proteins, most prominently IGF2BP3, and alter YTHDF2 recruitment to granules, including P-bodies and stress granules. Thus, base-editor screening reveals that individual residues within the YTHDF2 IDR are sufficient to couple protein-protein interactions, condensate localization, and the efficiency of m^6^A-dependent RNA degradation.

## RESULTS

### CRISPR base editor screens identify regulatory residues in YTHDF2

To comprehensively determine amino-acid residues that regulate YTHDF protein function in cells, we performed CRISPR base-editing screens with cytosine and adenine base editors (CBE and ABE) and sgRNAs tiled across the YTHDF2 coding sequence (Figure 1A). These editors introduce programmable C>T and A>G substitutions without requiring double-strand breaks and, when combined with Cas9-NG for relaxed PAM recognition, enable dense mutagenesis of protein-coding sequences ^27–30^. This strategy allows us to test the functional contribution of individual YTHDF2 residues in their native cellular context.

**Figure 1:**
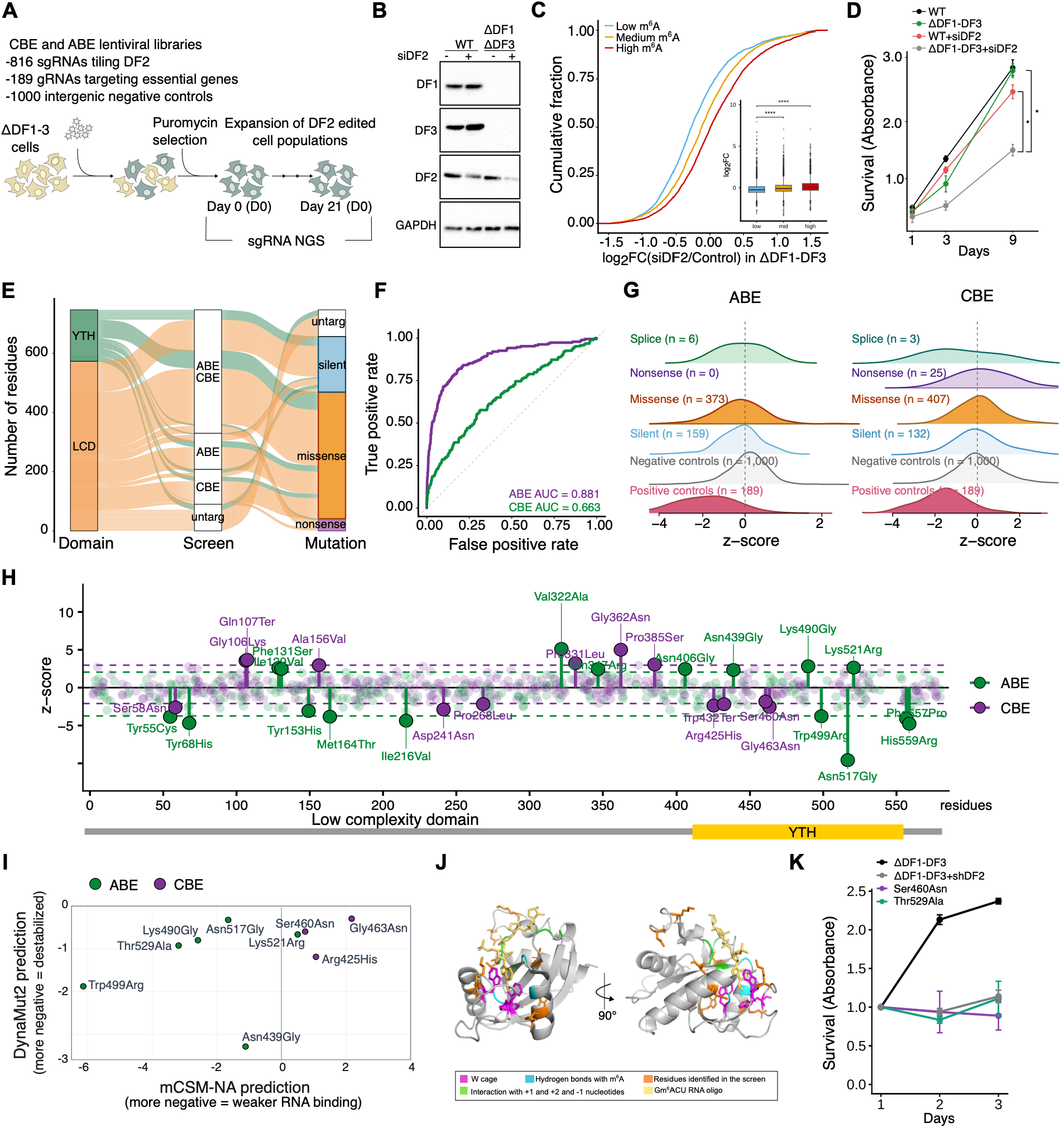
CRISPR base-editor screens identify functional residues in YTHDF2. **(A)** Schematic of the cytosine and adenine base-editor (CBE/ABE, respectively) screen strategy in ΔYTHDF1/ΔDF3 (ΔDF1-DF3) cells: lentiviral sgRNA libraries containing 816 sgRNAs tiling *YTHDF2*, 189 sgRNAs targeting essential genes, and 1000 intergenic non-targeting controls were cloned into all-in-one plasmids expressing either the CBE or the ABE base editor. Libraries were then transduced in ΔDF1/ΔDF3 cells, selected with puromycin, and sampled by sgRNA sequencing at Day 0 and Day 21. **(B)** Validation of YTHDF1 and YTHDF3 knockout and YTHDF2 depletion in HeLa cells. Immunoblot analysis confirmed loss of YTHDF1 and YTHDF3 in ΔDF1/ΔDF3 (ΔDF1-DF3) HeLa using the indicated YTHDF paralog-specific antibody. HeLa cells were also transfected with control or *YTHDF2*-targeting siRNAs, and YTHDF2 depletion was assessed 5 days after transfection. GAPDH was used as a loading control. **(C)** m^6^A mRNAs show an increase in expression upon depletion of YTHDF2 in ΔYTHDF1/ΔDF3 (ΔDF1-DF3) cells compared to control ΔDF1-DF3 cells. Transcripts were stratified into low, medium, and high m^6^A stoichiometry bins based on HeLa GLORI m^6^A measurements^2^. Bins correspond to low (<34% stoichiometry n=1039), mid (34-48% stoichiometry, n=646), and high (>48% stoichiometry n=4780) m^6^A stoichiometry. Cumulative distribution of log2 Fold Change (FC) in mRNA abundance after YTHDF2 depletion relative to control ΔDF1/ΔDF3 cells. Boxplots summarize the expression distribution for each m^6^A-site bin. Only significant p-values are shown. Two-tailed Mann-Whitney test; n=2, *p < 0.05, **p < 0.01, ***p < 0.001, ****p < 0.0001. **(D)** YTHDF2 depletion reduces proliferation in ΔYTHDF1/ΔDF3 (ΔDF1-DF3) cells. WST-1 proliferation assay of HeLa wild-type (WT), ΔDF1-3 cells, and upon YTHDF2 depletion (WT+siDF2, and ΔDF1-3+siDF2 cells) over 9 days. Data are mean ± SEM from n = 3 biological replicates. Wilcoxon rank-sum test; *p < 0.05, **p < 0.01, ***p < 0.001, ****p < 0.0001. **(E)** Design and coverage of the YTHDF2 base-editor sgRNA library. Diagram illustrating the YTHDF2 base-editor sgRNA library: number of residues targeted by domain (YTH, IDR), by screen (adenine base editor, ABE, cytidine base editor, CBE, ABE-CBE overlap, untargeted), and by predicted mutation outcome (missense, silent, nonsense), showing that the highest number of residues is covered by at least one of the two screens. Individual data are also presented in Figure S1A-B. **(F)** Receiver operating characteristic (ROC) curves evaluating the ability of guide-level robust z-scores to discriminate positive from negative control sgRNAs in the ABE and CBE screens. The ABE showed stronger classification performance than the CBE screen. **(G)** Distribution of guide-level z-scores for sgRNAs predicted to cause splice-site disruption, nonsense, missense, or silent mutations, and for negative and positive control sgRNAs, in the ABE (left) and CBE (right) screens. The number of gRNAs per category is indicated. **(H)** Residue-level robust z-scores across the YTHDF2 intrinsically disordered region (residues 1-409) and YTH domain (residues 410-544) for the ABE (green) and CBE (purple) screens at Day 21 relative to Day 0. Labeled residues denote top hits in each screen; yellow shading indicates the YTH domain. **(I)** Predicted effect of top YTH-domain hits on RNA binding and domain stability. mCSM-NA was used to estimate effects on m^6^A-RNA binding and DynaMut2 to estimate effects on YTH-domain stability. More negative mCSM-NA values indicate weaker predicted m^6^A-RNA binding; more negative DynaMut2 values indicate predicted destabilization. **(J)** Structural mapping of top YTH-domain screen hits onto the YTHDF2 YTH domain bound to m^6^A-containing RNA, highlighting the aromatic cage, hydrogen bonds with m^6^A, residues contacting flanking nucleotides, and residues identified by the screen. Screen-identified residues cluster in close spatial proximity to the bound methylated RNA, consistent with a role in supporting or fine-tuning m^6^A recognition. **(K)** Validation of selected YTH-domain hits by proliferation assay. WST-1 proliferation assay of ΔYTHDF1/ΔDF3 (ΔDF1-3), ΔDF1-3+shYTHDF2, and cells expressing individual sgRNAs targeting YTH-domain hits (Ser460Asn, Thr529Ala) over 3 days, normalized to Day 1. Data are mean ± SEM from n = 2 biological replicates.

Because YTHDF1, YTHDF2, and YTHDF3 act redundantly to promote m^6^A-mRNA decay ^8,9,11^, we used CRISPR to knock out YTHDF1 and YTHDF3 expression, allowing us to eliminate paralog compensation and thereby maximize the sensitivity of the screen to mutations in YTHDF2. We generated a YTHDF1/DF3 CRISPR double knockout HeLa cell line (ΔDF1/ΔDF3), leaving YTHDF2 (DF2) as the sole expressing YTHDF paralog (Figure 1B).

We further confirmed that YTHDF2 remained functionally competent to mediate m^6^A-dependent mRNA decay in this genetic background. To define m^6^A-marked transcripts, we used GLORI-based m^6^A maps generated in HeLa cells and stratified transcripts according to their m^6^A stoichiometry ^2^. Acute YTHDF2 knockdown through siRNAs confirmed that YTHDF2 depletion induced a selective increase in m^6^A-marked mRNA abundance relative to the ΔDF1/ΔDF3 background. As expected, the magnitude of the increase correlated with m^6^A stoichiometry, with more highly methylated transcripts showing stronger accumulation upon YTHDF2 depletion (Figure 1C; S1E). Moreover, YTHDF2 depletion reduced cellular viability compared to ΔDF1/ΔDF3 cells, confirming that YTHDF2 becomes essential for cellular fitness in the absence of YTHDF1 and YTHDF3 ^9,11^. (Figure 1D). Thus, the ΔDF1/ΔDF3 cell line provides a valuable system in which the functional consequences of sequence-level perturbations in YTHDF2 can be directly assessed at the levels of m^6^A-dependent mRNA regulation and cellular fitness.

To comprehensively interrogate YTHDF2 coding sequence function, we designed a lentiviral sgRNA library targeting all NG PAMs across the YTHDF2 open reading frame. The library contained 816 sgRNAs predicted to introduce base-editor-accessible substitutions across 490 amino acids, covering 85.93% of the YTH domain and 84.84% of the intrinsically disordered region (Figure 1E; S1A; S1B). In the ABE and CBE screens, 45.7% and 49.9% of guides were predicted to introduce missense substitutions, respectively. Silent substitutions were predicted for 25.7% of AM14 guides and 23.4% of CBE guides. In the CBE screen, 7.2% of guides were predicted to introduce nonsense substitutions (Figure 1E; S1B). As positive controls, we included 6 (ABE) and 3 (CBE) sgRNAs targeting genomic sites expected to disrupt YTHDF2 protein function, such as splice-proximal regions, together with sgRNAs directed against previously validated sites in common essential genes whose perturbation impairs cellular fitness. We further included 1,000 non-targeting sgRNAs as negative controls to define the background effects associated with sgRNA delivery, selection, and the screening workflow (Table S1). The library was cloned into lentiviral vectors that express the sgRNAs and contain either ABE8e fused to SpCas9-NG (ABE) or APOBEC1 fused to SpCas9-NG (CBE) ^30^ (see Methods).

To minimize the occurrence of multiple sgRNA integrations within individual cells while maintaining high representation of each library element at the population level, we transduced cells with the pooled lentiviral sgRNA library at a low multiplicity of infection (MOI < 0.3), thereby favoring a single perturbation per transduced cell. Cells were maintained for 21 days, and sgRNA abundance was quantified at Day 0 and Day 21 by deep sequencing in duplicate (Figure S1C).

To evaluate screen performance, we first analyzed guide-level behavior of internal positive and negative controls. For each screen, sgRNA abundance at Day 21 was compared to baseline at Day 0 to derive fold-change values using MAGeCK. Guide-level effects were then normalized relative to the negative-control distribution using a robust z-score. This transformation recentered the negative-control background around zero while mitigating baseline shifts between the ABE and CBE screens. After normalization, guide robust z-scores separated positive and negative controls with an AUC of 0.881 in ABE and 0.663 in CBE (Figure 1F), indicating the robustness of the screen and its ability to identify functional regulatory sites.

To identify YTHDF2 candidate regulatory residues, we next collapsed guide-level effects into residue-level scores. Each sgRNA was mapped to its predicted edited amino acid position, and robust ranking aggregation was used to obtain scores (z-scores) from all guides targeting the same residue within each screen. The resulting scores provided a quantitative measure of residue-specific selection and enabled the systematic prioritization of candidate functional sites across the YTHDF2 protein.

We next assessed whether perturbations predicted to compromise YTHDF2 function were depleted during the screen. sgRNAs targeting splice-proximal and intronic regions expected to disrupt YTHDF2 transcript processing exhibited negative selection by Day 21, consistent with reduced YTHDF2 activity conferring a fitness disadvantage in the ΔDF1/ΔDF3 background. (Figure 1G). Of note, consistent with the sensitized ΔDF1/ΔDF3 background, edits predicted to introduce premature termination codons, including Gln107Ter were also recovered by the screen at Day 21 (Figure 1H). These truncating alleles support the ability of the screen to detect disruptive YTHDF2 variants when compensation by YTHDF1 and YTHDF3 is removed.

As an additional internal validation of the screen, we also assessed whether the highest-scoring residues within the YTH domain corresponded to established determinants of m^6^A recognition. We identified 10 residues, 4 from the ABE screen and 6 from the CBE screen. Of these, 7 had a negative z-score, and 3 a positive z-score. Among the strongest negatively selected sites, we recovered Trp432, a conserved aromatic cage residue required for m^6^A binding (Figure 1H; S1D). We then used DynaMut2 ^31^ and mCSM-NA ^32^ to estimate whether the identified YTH-domain hits were predicted to alter domain stability or RNA binding, respectively (Figure 1I). Substitutions of many of the identified residues, including Thr529Arg and Trp499Ala, were predicted to destabilize both YTH-domain folding and RNA-binding potential as they mapped near the m^6^A-containing RNA-binding surface (Figure 1J). By contrast, substitutions at Ser460 or Arg425His were predicted to destabilize only the YTH fold. Taken together, the recovery of established m^6^A-contacting residues provides an internal validation of the screening strategy and supports its ability to resolve functionally relevant positions within the YTH domain with the sensitized ΔDF1/ΔDF3 background. The analysis further identified additional candidate residues predicted to influence RNA binding or domain stability.

To experimentally validate YTH-domain hits identified by the screen, we individually cloned selected sgRNAs and tested their effects on cell proliferation in ΔDF1/ΔDF3 cells. Cells expressing sgRNAs targeting candidate Ser460Asn and Thr529Ala YTH-domain residues showed reduced proliferation compared with cells expressing control sgRNAs over a three-day time course (Figure 1K).

Together, these results establish CRISPR base-editor screening as an effective strategy to map functional residues in YTHDF proteins. By combining a ΔDF1/ΔDF3 background with comprehensive mutagenesis of YTHDF2, guide-/and residue-level scoring, and structure-informed interpretation, this approach identifies both known m^6^A-recognition residues and additional regulatory sites required for YTHDF2 function.

### CRISPR base editor screens identify regulatory residues in YTHDF2 intrinsically disordered region

Notably, 67% of the highest-scoring functional residues localized outside the YTH domain, predominantly within the N-terminal intrinsically disordered region (IDR) of YTHDF2 (Figure 1H; S1D). Given the established involvement of YTHDF intrinsically disordered regions in protein-protein interactions, condensate partitioning, and phase-separation-associated behavior ^11,17,22^, we hypothesized that these screen-identified residues may modulate YTHDF2 by altering the IDR’s capacity for protein-protein interactions or condensate partitioning.

The screen identified 21 high-confidence IDR residues, defined as residues targeted by guides with robust z-scores beyond the screen-specific thresholds. Twelve residues were from the ABE screen and nine from the CBE screen. To investigate residue-specific regulation within the IDR, we focused subsequent analyses on high-confidence guides predicted to generate missense substitutions. We selected representative IDR-associated hits for individual validation. Cells were transduced with individual sgRNAs and the base editor, and puromycin-selected. Proliferation was monitored and quantified over 72 hours. Candidates that exhibited positive selection in the pooled screens, Gly362Asn and Val322Ala, did not confer reproducible proliferative advantages when evaluated individually (Figure S2A). By contrast, editing of Tyr55 (Tyr55Cys), Tyr68 (Tyr68His, Tyr68Cys), Thr87 (Thr87Ile), and Tyr153 (Tyr153His) reproducibly reduced proliferation relative to control lines, consistent with their depletion in the pooled screens (Figure 2A). These orthogonal validation experiments confirm that perturbation of specific residues within the IDR impairs cellular fitness.

**Figure 2:**
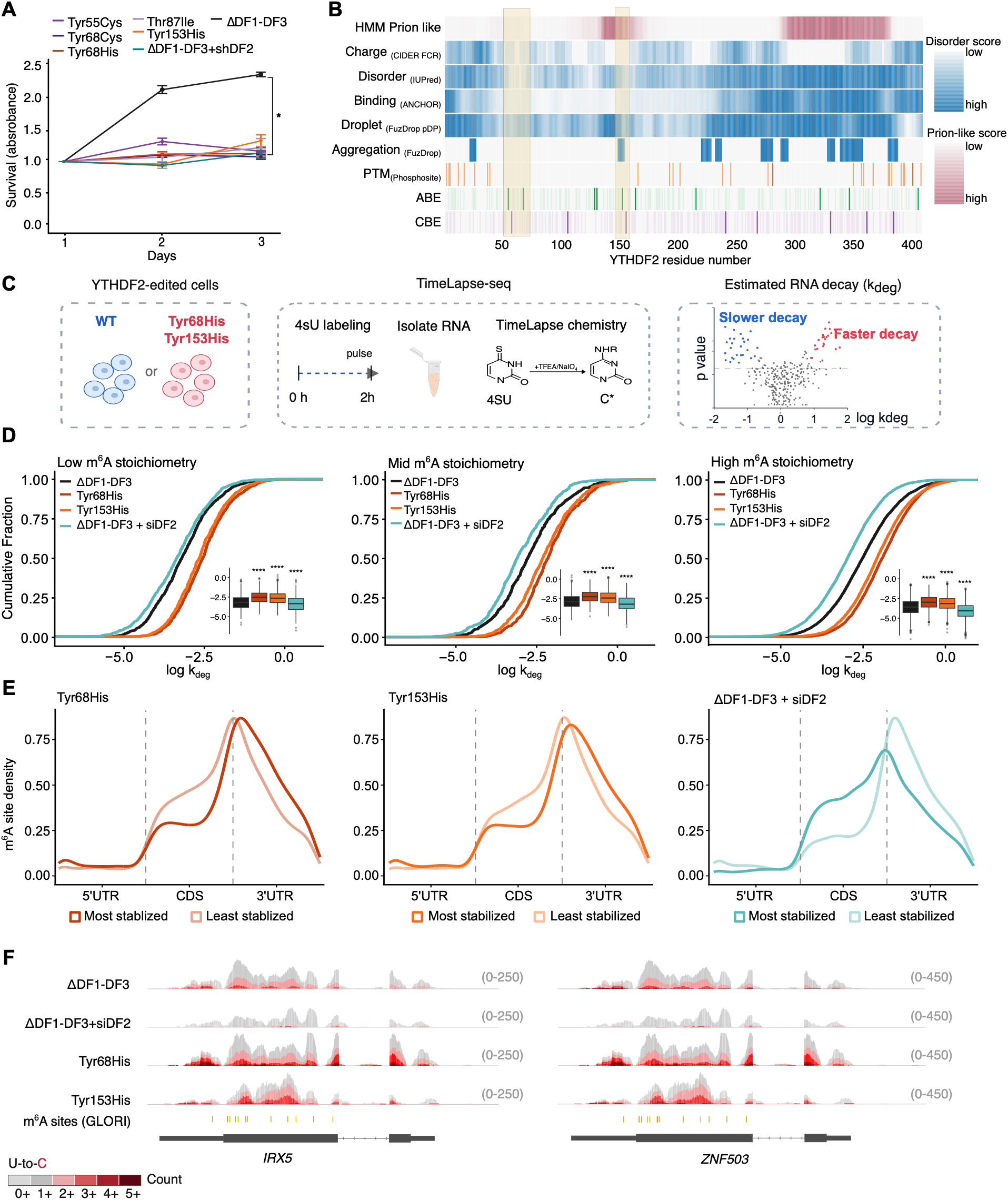
Base editor screens identify hyperactive YTHDF2 variants that enhance m^6^A-dependent mRNA decay. **(A)** Validation of candidate IDR residues identified by the base-editor screens. Proliferation of ΔYTHDF1/ΔDF3 HeLa cells, ΔYTHDF1-3+shYTHDF2 and cells expressing individual base-editor sgRNAs targeting identified IDR residues, measured by WST-1 proliferation assay over 3 days and relative to ΔYTHDF1/ΔDF3 (ΔDF1-DF3) control. All tested gRNAs predicted to target Tyr55Cys (p.adj = 0.036), Tyr68Cys (p.adj = 0.036), Tyr68His (p.adj = 0.0045), Thr87Ile (p.adj = 0.036), and Tyr153His (p.adj = 0.036) reduced proliferation relative to ΔYTHDF1/ΔDF3 control at Day 3. Data are mean ± SEM. Tyr68His (n = 4), Tyr153His (n = 3), Tyr55Cys (n = 2), Tyr68Cys (n = 2), and Thr87Ile (n = 2). Wilcoxon rank-sum test, BH-adjusted; star in Figure (*) indicates that all differences in proliferation are statistically significant as indicated in the legend. **(B)** Sequence-feature and screen-overlay map of the YTHDF2 N-terminal intrinsically disordered region (IDR, residues 1-409). Tracks (top to bottom): HMM prion-like score (PLAAC); charge patterning (CIDER FCR); predicted disorder (IUPred); predicted binding propensity (ANCHOR); droplet-promoting propensity (FuzDrop pDP); predicted aggregation propensity (FuzDrop); annotated post-translational modifications (PhosphoSitePlus). The bottom two tracks show positions assigned to individual guide RNAs in the ABE and CBE screens. Both positive and negative hits are shown in dark green for ABE and dark purple for CBE; hit direction is not encoded by color. Targeted, non-hit positions are shown with low transparency. **(C)** Experimental workflow for TimeLapse-seq analysis in YTHDF2 wild-type and edited cell lines. Tyr68His- and Tyr153His-edited ΔYTHDF1/ΔDF3 (ΔDF1-DF3) cells were generated by transduction with the corresponding base-editor sgRNA constructs, followed by puromycin selection and clonal isolation. Wild-type (WT) or Tyr68His/Tyr153His-YTHDF2 edited cells in the ΔDF1-DF3 background were labeled with 500uM 4-thiouridine (4sU) for 2 hours, RNA was isolated, and 4sU-containing nucleotides were chemically converted (NaIO₄) prior to sequencing to estimate transcriptome-wide RNA degradation rates (*k_deg*). As a control, non-NaIO4 converted RNA from the same 4sU-labeled samples was sequenced and used to estimate background signal for the *k_deg* calculation. **(D)** Tyr68His and Tyr153His increase degradation rates of m^6^A-modified transcripts. Cumulative distributions of log(kdeg) are shown for transcripts stratified into low, medium, and high m^6^A stoichiometry bins based on HeLa GLORI m^6^A measurements ^2^. Bins correspond to low (<34% stoichiometry n=1039), mid (34-48% stoichiometry, n=646), and high (>48% stoichiometry n=4780) m^6^A stoichiometry. Boxplots summarize the expression distribution for each m^6^A-site bin. Tyr68His and Tyr153His shift the *k_deg* distribution toward faster degradation rate relative to ΔYTHDF1/ΔDF3 (ΔDF1-DF3), with the magnitude of the shift increasing with m^6^A stoichiometry, whereas YTHDF2 depletion shifts the distribution toward slower degradation, consistent with stabilization of m^6^A-modified transcripts upon DF depletion. (Wilcoxon rank-sum test relative to ΔDF1/ΔDF3, ****p < 0.0001). **(E)** Metagene distribution of m^6^A site density across 5′UTR, CDS, and 3′UTR for the transcripts regulated by YTHDF2 IDR variants and upon YTHDF2 depletion. Genes were ranked based on the magnitude of change in k_deg relative to ΔDF1/ΔDF3 control in Tyr68His, Tyr153His and YTHDF2-depleted cells (ΔYTHDF1-DF3+siDF2). The “most” category has the 20% of m^6^A positive genes showing the largest change in *k_deg* (1264 genes per condition) and, the “least” group corresponds to the bottom 20% (1264 per condition). **(F)** Representative tracks of read coverage at *IRX5* and *ZNF503* in ΔDF1/ΔDF3, Tyr68His, Tyr153His, and ΔYTHDF1/ΔDF3+siYTHDF2 (ΔDF1-DF3+siDF2) cells. Read coverage is shown together with the 4sU-dependent U-to-C conversion signal, displayed in shades of red; darker red indicates higher conversion content. Increased U-to-C conversion content in Tyr68His and Tyr153His cells is consistent with faster RNA turnover and higher estimated degradation rates, whereas reduced conversion content in ΔDF1/ΔDF3+siDF2 cells is consistent with lower degradation rates and transcript stabilization upon YTHDF2 depletion. The distribution of the mapped m^6^A sites based on HeLa GLORI m^6^A measurements ^2^ is also indicated below.

We next asked whether these functional effects associated with these screen-identified IDR residues could be explained by primary-sequence features known to shape disordered protein behavior. We analyzed the YTHDF2 IDR using established sequence-prior metrics, including PLAAC/HMM.PrD-like scoring for prion-like composition ^33^, localCIDER-style analysis for charge patterning and IDR sequence grammar ^34^, IUPred/ANCHOR for disorder and protein-binding propensity (i.e. predicted capacity to mediate protein-protein interactions) ^35^, FuzDrop for droplet-promoting propensity ^36^ and annotated post-translational modification sites ^37^. Consistent with previous analyses of YTHDF2 IDR ^38,39^, these predictors show that the YTHDF2 IDR is not compositionally uniform but contains discrete sequence-defined subregions.

We then compared these sequence-prior annotations with the ABE and CBE residue-level screen profiles (Figure 2B). This comparison showed that a subset of functional IDR residues mapped to sequence-defined regions predicted to have prion-like or droplet-promoting character. Tyr153 mapped within a short region spanning residues 143-157. Similarly, additional screen hits overlapped a second HMM.PrD-like region in a distal IDR area (300-360). In contrast, Tyr68, along with Tyr55 and Thr87, showed strong negative z-scores in our screen despite not being annotated by any of the sequence-based priors used in our analysis. Tyr68 also showed the highest guide concordance across independent sgRNAs (average score -2.72; Tyr68His: -3.56; Tyr68His: -1.71; Tyr68Cys: -2.79; Table S1).

Collectively, these results indicate that the YTHDF2 IDR contains a residue-resolved functional architecture that is only partially captured by current sequence-based models of intrinsically disordered protein behavior.

### IDR mutations identify hyperactive YTHDF2 variants that enhance m^6^A-dependent mRNA decay

We selected Tyr68 and Tyr153 for mechanistic analysis because they represented two complementary classes of IDR regulatory residues. Tyr153 was a top-scoring residue within a region supported by sequence-based IDR predictors, whereas Tyr68 was a top-scoring functional residue that would not have been prioritized from IDR sequence features alone (Figure 2B). Notably, both Tyr68 and Tyr153 are conserved across YTHDF1, YTHDF2, and YTHDF3 (Figure S2B), suggesting that these positions may represent conserved regulatory residues within the YTHDF IDR family.

To investigate the mechanism underlying these phenotypes, we generated stable edited ΔDF1/ΔDF3 cell lines carrying Tyr68His or Tyr153His by transducing cells with the corresponding base-editor sgRNA constructs and isolating edited populations or clones for downstream assays (Figure 2C). Sanger sequencing confirmed the expected editing event at Tyr68 and Tyr153 in the edited lines (Figure S2C). Independently isolated edited clones showed consistent proliferation defects relative to unedited ΔDF1/ΔDF3 control cells, confirming that the impaired growth phenotype is a direct effect of the IDR mutations (Figure S2D, S2E-S2F). In the case of Tyr68His, two sgRNAs predicted to generate distinct amino-acid substitutions at Tyr68, Tyr68His and Tyr68Cys, were individually cloned and introduced into ΔDF1/ΔDF3 cells. Both sgRNAs reduced cellular fitness in validation assays (Figure 2A). Overall, these results suggest that the proliferation defect reflects a requirement for Tyr68 and Tyr153 rather than a guide-specific effect or a single-substitution-specific artifact. Of note, because these mutations caused progressive growth impairment under extended culture conditions, all experiments were performed within a maximum of 8 passage windows.

To determine whether the Tyr68His and Tyr153His substitutions alter YTHDF2-dependent, m^6^A-mediated mRNA decay, we quantified transcriptome-wide mRNA turnover using TimeLapse-seq. Newly synthesized transcripts were labeled with 4-thiouridine for 2 hours, which was subsequently recoded into a cytidine analog through oxidative-nucleophilic aromatic substitution, generating characteristic U-to-C conversions upon sequencing. After sequencing, we then used EZbakR ^40^, which models these U-to-C conversions to derive per-transcript degradation rate constants, *kdeg* (Figure 2C). Thus, the *kdeg* is a direct readout of turnover, where a *high kdeg* corresponds to a rapidly degraded, short-lived transcript, and a *lower kdeg* to a more stable one.

TimeLapse-seq libraries showed strong replicate concordance across ΔDF1/ΔDF3, Tyr68His, Tyr153His samples, as well as the corresponding unlabeled controls (Figure S2G). In addition, 4sU-dependent conversion rates were clearly separated from background conversion rates in all conditions (Figure S2H). These quality-control analyses confirmed consistent metabolic labeling, chemical conversion, and sequencing performance across samples, supporting comparison of *kdeg* estimates between genotypes.

We next examined whether the effects of YTHDF2 variants on mRNA turnover varied as a function of transcript methylation. To this end, transcript-specific degradation rates were integrated with a HeLa m^6^A map generated using GLORI, which selectively converts unmethylated adenosines through glyoxal- and nitrite-mediated deamination while leaving m^6^A-modified adenosines intact^2^. The resulting nucleotide-conversion pattern enables m^6^A sites to be identified at single-nucleotide resolution and their methylation stoichiometry to be quantified. Transcripts were subsequently stratified into low-, intermediate-, and high-methylation groups according to the mean m^6^A stoichiometry across sites assigned to each transcript. Across all three methylation groups, m^6^A-modified transcripts showed an overall 1.7-fold shift toward higher *kdeg* in both Tyr68 and Tyr153 mutant cells compared with wild-type YTHDF2 cells, indicating broadly enhanced degradation of methylated transcripts in the mutant lines. Although this effect was not strictly proportional to m^6^A stoichiometry, highly methylated YTHDF-sensitive transcripts are already rapidly degraded at baseline, which may limit the dynamic range for detecting further enhanced degradation. (Figure 2D-Table S2).

To establish a loss-of-function reference for YTHDF2-dependent mRNA decay, we compared the mutant profiles with transcriptome-wide mRNA turnover measurement in YTHDF2-depleted ΔDF1/ΔDF3 cells (Figure S2G, S2H). As expected, depletion of the remaining YTHDF paralog reduced *kdeg* for m^6^A-modified transcripts by 1.6-fold, consistent with stabilization of YTHDF-sensitive RNAs. This is the opposite of what we observed in Tyr68 and Tyr153-mutant cells. Thus, mutations in Tyr68 and Tyr153 act as hyperactive YTHDF2 variants that enhance m^6^A-dependent mRNA degradation.

We next determined whether Tyr68His and Tyr153His act on the same m^6^A-decay program normally controlled by YTHDF2. Recent studies have shown that YTHDF2-dependent mRNA decay is particularly associated with m^6^A sites located within coding sequences (CDS) ^41,42^. We therefore examined the metagene distribution of m^6^A sites on transcripts with the largest changes in degradation rate after YTHDF2 depletion and in the Tyr68His and Tyr153His mutants (Figure 2E, S2I). Consistent with previous findings, transcripts stabilized by YTHDF1-2-3 depletion showed the strongest enrichment for coding sequence-localized m^6^A sites. Notably, transcripts destabilized by the Tyr68His and Tyr153His variants exhibited a CDS-enriched distribution of m^6^A sites. Thus, although YTHDF2 depletion and the two variants exerted opposing effects on transcript stability, they converged on the same class of CDS-m^6^A-containing targets. For example, *IRX5* and *ZNF503* were stabilized upon YTHDF2 depletion, as reflected by a low proportion of U-to-C mutated reads despite constant overall coverage compared to control. In contrast, both transcripts were destabilized in the Tyr68His and Tyr153His mutant lines with a higher fraction of reads carrying U-to-C mutations, consistent with increased turnover. Both genes carry m^6^A sites predominantly within the CDS (*IRX5*: 80% of sites at 86.2% average stoichiometry; *ZNF503*: 92.9% of sites at 68.1% average stoichiometry) (Figure 2F). Thus, the same m^6^A-modified CDS-localized targets that are stabilized upon YTHDF2 depletion are destabilized by the Tyr68His and Tyr153His variants, confirming that these mutations enhance the canonical YTHDF-dependent decay program.

Collectively, these findings establish Tyr68 and Tyr153 as negative regulatory determinants of YTHDF2 activity. Mutation of either residue enhances the degradation of m^6^A-modified transcripts, revealing a previously unrecognized role for specific residues within the YTHDF2 intrinsically disordered region in controlling the magnitude of m^6^A-dependent mRNA decay.

### Hyperactive YTHDF2 variants control YTHDF2 protein interaction network

Having established that the Tyr68His and Tyr153His substitutions potentiate YTHDF2-mediated, m^6^A-dependent mRNA decay, we next sought to define the molecular mechanism underlying their gain-of-function effects. Since Tyr68 and Tyr153 lie outside the YTH domain, the hyperactive phenotype was unlikely to be explained by direct changes in m^6^A recognition. However, these substitutions could indirectly alter RNA recognition through long-range conformational coupling, change in intramolecular interactions, or altered accessibility of the YTH domain ^43,44^. We therefore tested the YTHDF2 mutant-RNA interaction experimentally by purifying recombinant wild-type, Tyr68His, and Tyr153His YTHDF2 proteins from *E.coli* and measuring binding to an m^6^A-containing RNA probe by Electromobility Shift Assay (EMSA) (Figure S3A; S3B). Both IDR mutants bound m^6^A-RNA comparably to wild-type YTHDF2, indicating that their increased decay activity is not caused by enhanced intrinsic m^6^A binding.

We next investigated whether Tyr68 and Tyr153 increase protein levels. Immunoblotting confirmed that YTHDF2 protein levels were comparable between wild-type, Tyr68His, and Tyr153His edited cells, indicating that the mutations do not alter YTHDF2 protein expression (Figure S2F). We next asked whether the Tyr68His and Tyr153His substitutions instead altered YTHDF2 protein turnover, which could indirectly increase YTHDF2 abundance over time without changing steady-state protein levels at a single time point ^45^. To evaluate protein stability levels, we blocked new protein synthesis with cycloheximide. Cells expressing wild-type YTHDF2,

Tyr68His, or Tyr153His YTHDF2 were treated with cycloheximide, and YTHDF2 protein levels were monitored and quantified over two hours. Within two hours, we observed a 40-50% reduction in wild-type YTHDF2 protein levels, as previously observed ^45^. Tyr68His and Tyr153His showed degradation kinetics comparable to wild-type YTHDF2 over this time course (Figure S3C). Thus, the enhanced decay activity of the IDR mutants cannot be explained by increased m^6^A-RNA binding or increased YTHDF2 protein stability.

Having established that the observed hyperactive phenotype is not mediated by significant changes in direct m^6^A-binding affinity or protein levels, we next investigated whether Tyr68 and Tyr153 instead regulate YTHDF2 through altered protein-protein interactions. Recently, several studies have established that intrinsically disordered regions in RNA-binding proteins function as modular interaction platforms that recruit cofactors and thereby influence RNA fate ^23,46^. We therefore hypothesized that Tyr68 and Tyr153 participate in protein-protein interactions that modulate YTHDF2 activity on m^6^A degradation. In this model, wild-type YTHDF2 associates with one or more inhibitory factors that inhibit m^6^A-dependent decay, whereas mutation of Tyr68 or Tyr153 disrupts this association and releases YTHDF2 into a hyperactive state.

We therefore sought to first define the endogenous YTHDF2-proximal proteome and then determine which of the interactors could have an inhibitory role on YTHDF2-mediated mRNA decay. Previous proximity-labeling and interaction-mapping studies have shown that YTHDF proteins associate with both mRNA decay factors ^8,18,19^ and RNA-binding proteins linked to transcript stabilization and condensate regulation ^47^. However, most of these studies relied on overexpressed tagged YTHDF proteins, making it unclear whether these candidate regulatory associations occur in native YTHDF2 cellular conditions. To address this, we adapted μMAP proximity labeling to endogenous YTHDF2 ^48^. In this approach, fixed and permeabilized cells are stained with a primary antibody against endogenous YTHDF2, followed by a secondary antibody conjugated to a photocatalyst. Upon light activation, proteins in proximity to endogenous YTHDF2 are biotinylated and subsequently enriched for mass-spectrometry analysis (Figure 3A). To increase confidence in the resulting interaction map and reduce fixation-specific artifacts, we performed endogenous YTHDF2 μMAP using two fixation conditions, paraformaldehyde and methanol, each benchmarked against an isotype-matched IgG control.

**Figure 3:**
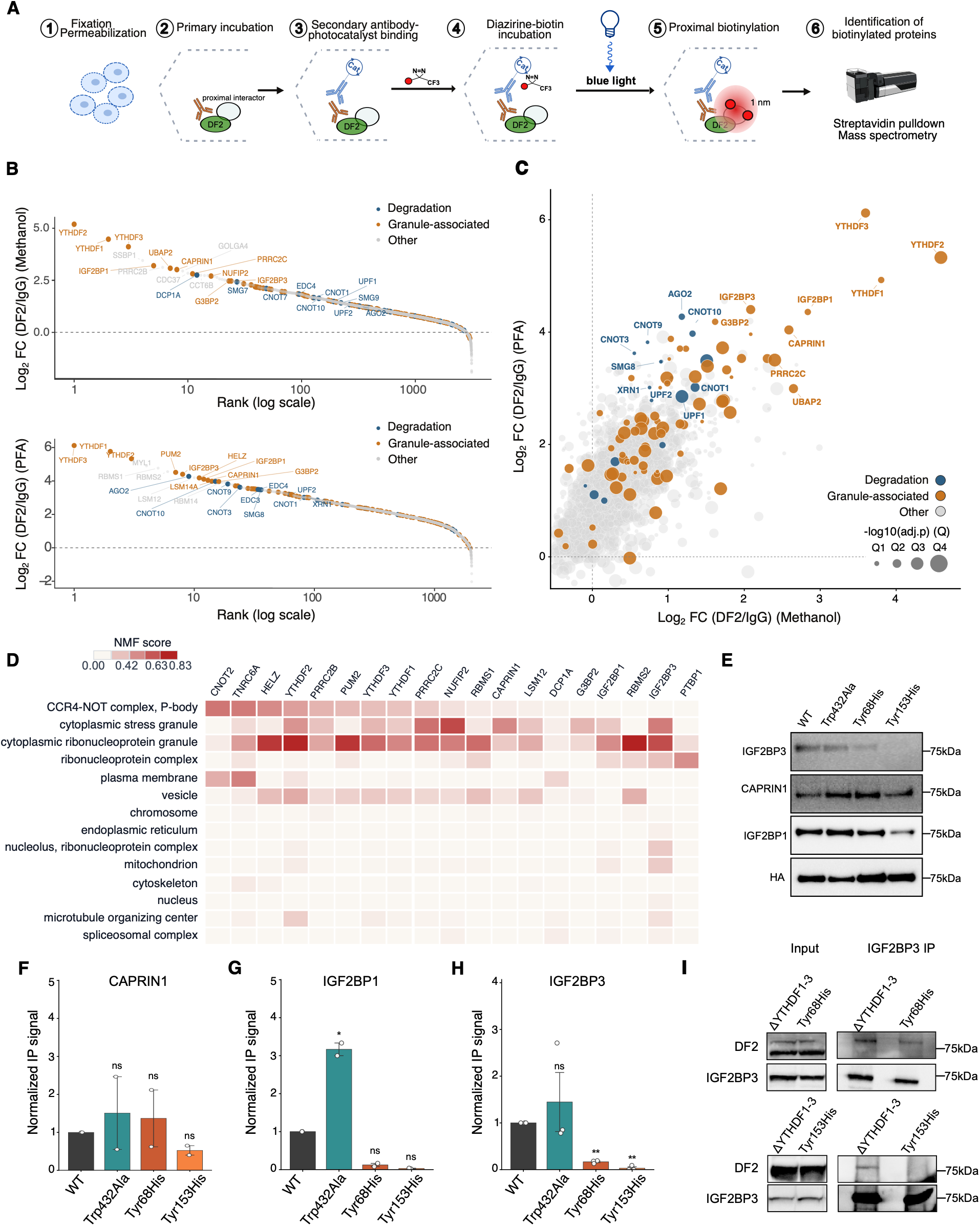
Hyperactive YTHDF2 variants control the YTHDF2 protein interaction network. (**A**) Schematic of YTHDF2 proximity labeling adapted from μMAP ^48^ to detect endogenous YTHDF2 interactome. Fixed and permeabilized HeLa cells were stained with a primary antibody against endogenous YTHDF2, followed by incubation with a photocatalyst-conjugated secondary antibody. Upon light activation in the presence of diazirine-biotin probe, proteins proximal to endogenous YTHDF2 are biotinylated within a short labeling radius and subsequently enriched for mass-spectrometry analysis. **(B)** Ranking of endogenous YTHDF2-proximal proteins identified by μMAP following methanol (top) or paraformaldehyde (bottom) fixation. All detected proteins were ranked by enrichment over the corresponding IgG control, calculated as log2 fold change. Significantly enriched proteins are annotated and color-coded according to reported functions in RNA degradation (blue) or RNA granules (orange); proteins without these annotations are shown in gray. **(C)** Endogenous YTHDF2 μMAP identifies reproducible YTHDF2-proximal proteins across fixation conditions. Scatter plot comparing protein enrichment over IgG control in paraformaldehyde-fixed (y-axis, log2 fold-change) versus methanol-fixed (x-axis, log2 fold-change) cells. Proteins are colored by functional category based on GO annotation (Degradation, blue; Granules, orange; Other, gray). Point size reflects the significance of enrichment, divided into quartiles based on −log10(adjusted P-value) across the two fixation conditions (Q1, least significant; Q4, most significant). 137 proteins enriched in both conditions are plotted and prioritized as high-confidence YTHDF2-proximal interactors. **(D)** Non-negative matrix factorization (NMF)-based annotation of the top endogenous YTHDF2 μMAP interactors. Heatmap showing the localization profiles of the top YTHDF2-proximal proteins annotated using the NMF subcellular localization framework from Youn et al. ^51^. Rows indicate RNA-protein/organelle-associated modules; columns indicate individual YTHDF2-proximal proteins; color indicates NMF localization score (scale, 0.00-0.83). Interactors have been ranked by their similarity to the CNOT2 interactome (first column). This analysis highlights YTHDF2 and TNRC6A as proteins with interactors similar to those of CNOT2. IGF2BPs’ interactors have low similarity to CNOT interactors. **(E)** Proximity biotinylation assay to test the effect of IDR variants on candidate YTHDF2 interactors. ΔDF1/ΔDF3 cells were transiently transfected with BirA-tagged YTHDF2-HA constructs encoding wild-type YTHDF2 (WT), the m^6^A-binding-deficient control mutant Trp432Ala, or Tyr68His or Tyr153His YTHDF2. Biotinylated proteins were enriched by streptavidin pulldown and immunoblotted for IGF2BP3, CAPRIN1, and IGF2BP1. HA immunoblotting was used to assess the immunoprecipitation efficiency across conditions together with the input shown in Figure S3. **(F–H)** Quantification of streptavidin pulldown signal for CAPRIN1 (**F**), IGF2BP1 (**G**), and IGF2BP3 (**H**) from (**E**), normalized to BirA-tagged YTHDF2-HA expression shown in Figure S3. Data are mean ± SEM from n = 2-3 biological replicates as indicated. T test; ns, non-significant, *p < 0.05, **p < 0.01, ***p < 0.001, ****p < 0.0001. **(I)** Co-immunoprecipitation of endogenous IGF2BP3 from ΔDF1/ΔDF3 cells expressing endogenous wild-type YTHDF2, Tyr68His, or Tyr153His YTHDF2. IGF2BP3 was immunoprecipitated, and co-purification of YTHDF2 was assessed by immunoblotting. Compared to wild-type YTHDF2, both Tyr68His and Tyr153His showed 87.5% and almost complete depleted association with IGF2BP3, respectively, indicating that these IDR residues are required to maintain the YTHDF2-IGF2BP3 interaction. Input and IGF2BP3 immunoprecipitation fractions were immunoblotted for YTHDF2 and IGF2BP3.

To identify high-confidence endogenous YTHDF2-proximal proteins, we compared enrichment over IgG controls in paraformaldehyde- and methanol-fixed cells. To do so, we plotted each protein according to its log2 fold enrichment over IgG in the two conditions. We identified 1160 proteins as significantly enriched using paraformaldehyde and 154 proteins using methanol, consistent with the higher overall protein yield typically obtained with paraformaldehyde fixation^49^.

We then focused on the 137 proteins enriched under both fixation conditions as a high-confidence endogenous YTHDF2-proximal proteome (Figure 3C). This proximal proteome recovered proteins previously reported in YTHDF2 interaction datasets, supporting endogenous μMAP as a strategy to capture native YTHDF2-associated protein interactions (Figure S3D) ^47,48,50^. This set included YTHDF1 and YTHDF3, consistent with the known association among YTHDF paralogs ^9,11,39^. As expected from the established role of YTHDF2 in m^6^A-dependent mRNA turnover ^8,10^, the endogenous proximity proteome was also enriched for factors associated with RNA decay, including CNOT complex proteins and TNRC6A. Importantly, it also recovered RNA-binding proteins linked to transcript stabilization and ribonucleoprotein granule biology, including IGF2BP1, IGF2BP3, and CAPRIN1, among the most significant and reproducibly enriched proteins (Figure 3C). Because these factors were recovered across two fixation conditions using endogenous YTHDF2 labeling, these data support their presence in the native YTHDF2 protein neighborhood rather than reflecting artifacts of tagged YTHDF overexpression. Thus, endogenous YTHDF2 μMAP captures both the expected decay-associated YTHDF2 interactors and a robust native proximity signal composed of RNA-stabilizing and granule-associated proteins.

To determine whether these proteins represented a functional neighborhood distinct from canonical decay machinery, we next annotated the top μMAP-identified YTHDF2-proximal proteins using the Youn et al. localization framework ^51^. This framework uses proximity-labeling profiles of RNA-binding and RNA-regulatory proteins to assign proteins to modules corresponding to distinct subcellular compartments and ribonucleoprotein assemblies (NMF, non-negative matrix factorization). We ordered the YTHDF2-proximal proteins by similarity to the CNOT2 profile, using CNOT2 as a reference for decay-associated/P-body machinery (Figure 3D). This analysis positioned YTHDF2, TNRC6A, and CNOT-associated proteins near the CNOT-like profile, consistent with their roles in mRNA repression and decay. In contrast, IGF2BP1, IGF2BP3, CAPRIN1, PUM2, PRRC2B/C, HELZ, and LSM12 mapped away from this profile and aligned with cytoplasmic RNP granule or stress-granule-associated modules. Thus, the endogenous YTHDF2-proximal proteome contains both canonical decay-associated factors and a distinct RNA-stabilizing/granule-associated protein neighborhood.

Among the proteins most distant from the CNOT-like decay profile, IGF2BP1 and IGF2BP3 were among the strongest and most reproducible YTHDF2-proximal proteins (Figure 3D). IGF2BP1, IGF2BP2, and IGF2BP3 are a family of proposed non-canonical m^6^A-reader proteins that, in contrast to YTHDF proteins, stabilize target transcripts and enhance their translation ^52,53^. We therefore hypothesized that association with IGF2BP proteins may attenuate YTHDF2-mediated mRNA degradation, and that disruption of this association by the Tyr68His and Tyr153His mutations could contribute to the hyperactive phenotypes.

To determine whether Tyr68 or Tyr153 are required for the association between YTHDF2 and IGF2BP proteins, we expressed BirA-tagged wild-type YTHDF2, Tyr68His, or Tyr153His and performed proximity labeling followed by immunoblotting for candidate interactors ^54^. Substitution of either Tyr68 or Tyr153 markedly reduced proximity labeling of IGF2BP1 and IGF2BP3, whereas labeling of other YTHDF2-proximal proteins, including CAPRIN1, was largely preserved (Figure 3E-H, S3G). Proximity to IGF2BP2, the third member of the IGF2BP family ^55^, was also reduced (Figure S3F, S3G), although the most prominent effect was observed for IGF2BP3, with no significant change in IGF2BP3 protein levels observed across conditions (Figure S3H). These results indicate that the two substitutions selectively perturb the proximity of YTHDF2 to IGF2BP family members, most prominently IGF2BP3, rather than broadly disrupting its proximal interaction network.

To determine whether YTHDF2 proximity to IGF2BP proteins requires YTHDF2 recognition of m^6^A-modified RNA, we included the m^6^A-binding-deficient Trp432Ala variant in the proximity-labeling experiments. In contrast to Tyr68His and Tyr153His, Trp432Ala did not reduce proximity labeling of IGF2BPs (Figure 3G-H, S3F-S3G). Thus, loss of m^6^A binding does not phenocopy the Tyr68His or Tyr153His variants, indicating that YTHDF2 proximity to IGF2BP proteins is not simply driven by co-occupancy of m^6^A-modified RNA substrates. Instead, these data support a model in which Tyr68 and Tyr153 define an IDR-dependent regulatory interface that promotes YTHDF2 proximity to IGF2BP proteins.

We further validated the YTHDF2-IGF2BP3 interaction by native co-immunoprecipitation of IGF2BP3 from cells expressing wild-type, Tyr68His, and Tyr153His YTHDF2. Although YTHDF2 recovery was low under native lysis conditions, IGF2BP3 co-purified with wild-type YTHDF2, whereas its association with both IDR variants was substantially reduced (Figure 3I). These results confirm the proximity-labeling data (Figure 3E-H).

Together, these findings identify Tyr68 and Tyr153 as important determinants of the YTHDF2-IGF2BP3 interaction and support a model in which disruption of this protein-protein interaction contributes to the enhanced m^6^A-dependent decay activity of the two variants. Thus, two of the IDR residues identified by the screen control YTHDF2 through a potential inhibitory protein-protein interaction that limits YTHDF2 function.

### Hyperactive YTHDF2 variants control YTHDF2 partitioning into cytoplasmic RNA granules

IDRs provide interaction valency that can promote protein partitioning into biomolecular condensates and contribute to the assembly of cytoplasmic RNA granules ^22^. Similarly, the IDRs of YTHDF proteins have been shown to promote their localization to both P-bodies and stress granules, with multivalent m^6^A-modified RNAs acting as scaffolds for YTHDF protein recruitment to condensates ^17,56^. We therefore examined whether Tyr68His and Tyr153His alter either the formation of these compartments or the recruitment of YTHDF2 into these compartments.

We first asked whether YTHDF2 itself was required for P-body maintenance in a ΔDF1/DF3 background. To acutely deplete the remaining YTHDF paralog while minimizing secondary adaptations, we generated an inducible shRNA system that reduced YTHDF2 abundance by 74% within 24 hours (Figure S4A). We then quantified P-bodies using EDC4, a core P-body component and established P-body marker ^57^. Acute YTHDF2 depletion did not significantly alter the number of EDC4-positive P-bodies per cell (Figure S4B). Thus, in ΔDF1/DF3 cells, loss of the remaining YTHDF protein is not sufficient to alter P-body number.

We next asked whether the IDR variants altered the partitioning of YTHDF2 into P-bodies. Because P-bodies have been proposed to provide sites where YTHDF proteins engage m^6^A-modified transcripts for degradation ^16^, we reasoned that hyperactive YTHDF2 mutants might show increased localization to P-bodies. To test this, we calculated a recruitment score by measuring the background-corrected YTHDF2 fluorescence within EDC4-positive compartments and normalizing it to the total YTHDF2 fluorescence in the same cell (Figure 4A). Tyr68His and Tyr153His increased the fraction of YTHDF2 localized to P-bodies by 27% and 22%, respectively, relative to the parental ΔDF1/ΔDF3 cells in independently analyzed clones (Figure 4B; S4C). Neither variant altered the number of EDC4-positive P-bodies (Figure 4C; S4D). Thus, the hyperactive IDR variants do not increase P-body number but instead enhance YTHDF2 partitioning into pre-existing decay-associated P-bodies.

**Figure 4:**
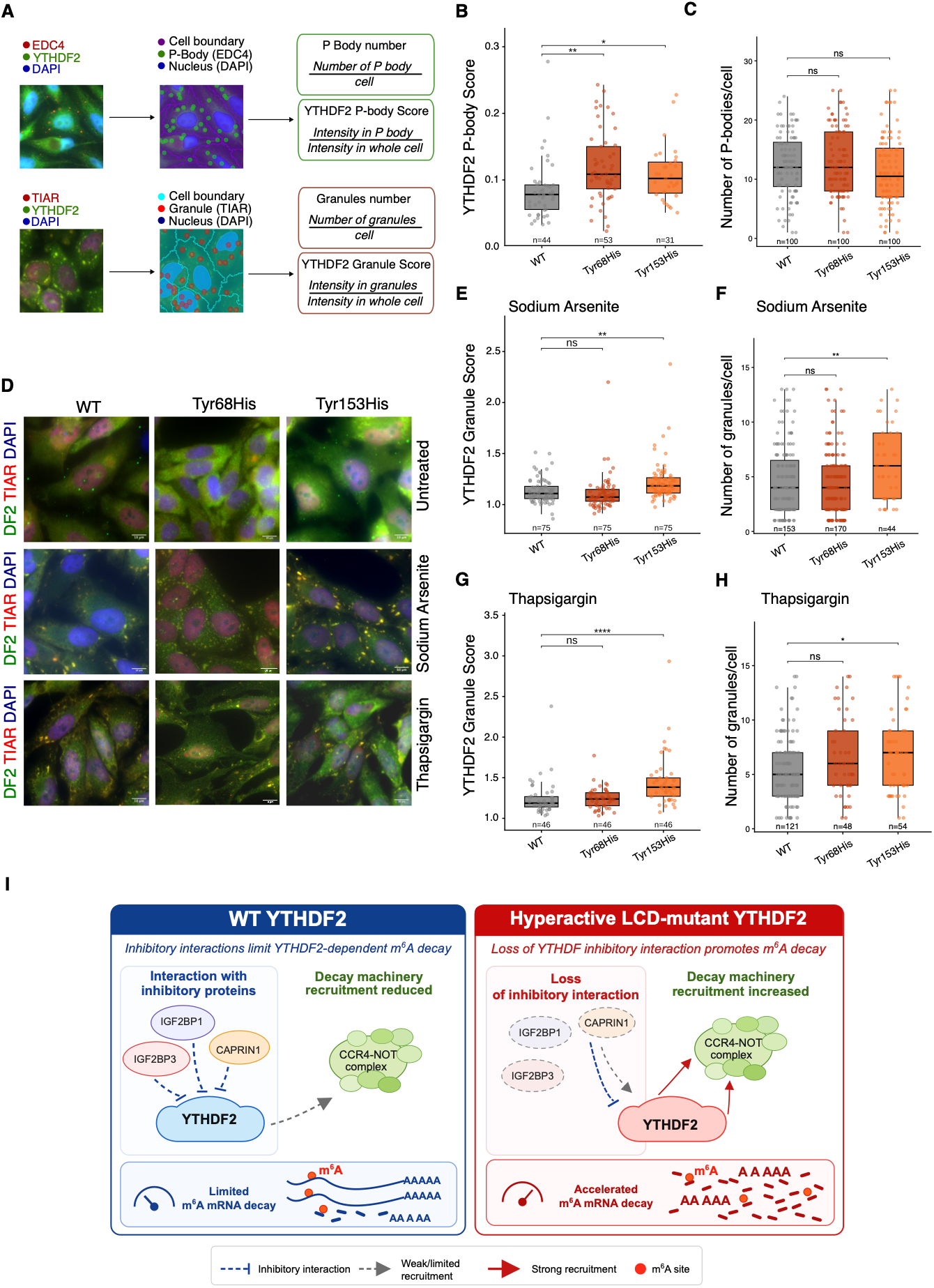
Hyperactive YTHDF2 variants control YTHDF2 localization to P-bodies and stress granules. **(A)** Schematic illustrating the YTHDF2 condensate-partitioning analysis workflow. P-bodies or stress granules were identified by segmenting EDC4- and TIAR-positive puncta, respectively, from the surrounding cytoplasm using Nikon NIS-elements GA3 software. The YTHDF2 recruitment score was calculated as the background-corrected integrated YTHDF2 fluorescence within the segmented compartment divided by YTHDF2 fluorescence in the corresponding cell. **(B)** YTHDF2 recruitment to EDC4-positive P-bodies in ΔDF1/ΔDF3, Tyr68His, and Tyr153His cells, quantified as described in (**A**). Tyr68His and Tyr153His increased YTHDF2 recruitment by 27.0% and 21.9%, respectively, relative to ΔDF1/ΔDF3 cells. Wilcoxon rank-sum test, BH-adjusted, *p < 0.05, **p < 0.01, ***p < 0.001, ****p < 0.0001, (n = 44 ΔDF1-DF3, 53 Tyr68His, 31 Tyr153His cells from 2 independent experiments). **(C)** Number of P-bodies per cell in ΔDF1/ΔDF3, Tyr68His, and Tyr153His cells. Neither Tyr68His nor Tyr153His significantly altered P-body number relative to ΔDF1/ΔDF3. Wilcoxon rank-sum test, BH-adjusted; ns = non-significant (n = 100 ΔDF1-DF3, 100 Tyr68His, 100 Tyr153His cells). **(D)** Representative immunofluorescence images of YTHDF2 (green), TIAR (red), and nuclei stained with DAPI (blue) in ΔDF1/ΔDF3, Tyr68His, and Tyr153His cells, under unstressed conditions or following treatment with sodium arsenite or thapsigargin to induce oxidative or endoplasmic reticulum stress, respectively. Scale bar: 10 µm. **(E)** YTHDF2 enrichment at stress granules under oxidative stress (sodium arsenite), quantified as described in (**A**). Under oxidative stress, only Tyr153His significantly increased YTHDF2 recruitment to stress granules relative to ΔDF1/ΔDF3; Tyr68His showed no significant change. Wilcoxon rank-sum test vs. ΔDF1/ΔDF3, BH-adjusted. *p < 0.05, **p < 0.01, ***p < 0.001, ****p < 0.0001 (n = 75 ΔDF1/ΔDF3, 75 Tyr68His, 75 Tyr153His cells from 2 independent experiments). (**F**) Stress-granule number per cell following oxidative stress (Sodium Arsenite). Tyr153His significantly increased stress-granule number, whereas Tyr68His did not significantly alter stress-granule abundance. Wilcoxon rank-sum test vs. ΔDF1/ΔDF3, BH-adjusted. *p < 0.05, **p < 0.01, ***p < 0.001, ****p < 0.0001 (n = 153 ΔDF1/ΔDF3, 170 Tyr68His, 44 Tyr153His cells from 2 independent experiments). (**G**) YTHDF2 enrichment at stress granules under endoplasmic reticulum stress (Thapsigargin), quantified as described in (**A**). Tyr153His significantly increased YTHDF2 recruitment to stress granules relative to ΔDF1/ΔDF3, whereas Tyr68His showed no significant change. Wilcoxon rank-sum test vs. ΔDF1/ΔDF3, BH-adjusted. *p < 0.05, **p < 0.01, ***p < 0.001, **** p < 0.0001 (n = 46 ΔDF1/ΔDF3, 46 Tyr68His, 46 Tyr153His cells from 2 independent experiments). (**H**) Stress-granule number per cell following endoplasmic reticulum stress (Thapsigargin). Tyr153His significantly increased stress-granule number, whereas Tyr68His did not significantly alter stress-granule abundance. Wilcoxon rank-sum test vs. ΔDF1/ΔDF3, BH-adjusted. *p < 0.05, **p < 0.01, ***p < 0.001, ****p < 0.0001 (n = 121 ΔDF1/ΔDF3, 48 Tyr68His, 54 Tyr153His cells from 2 independent experiments). (**I**) Proposed model showing IDR-mediated YTHDF2 regulation. In wild-type cells, interactions between the YTHDF2 IDR and RNA-stabilizing or granule-associated proteins, including IGF2BP3, may inhibit m^6^A-mRNA decay. Mutations of regulatory IDR residues alter these interactions, enhancing m^6^A-dependent mRNA decay.

We next examined whether the variants similarly affected YTHDF2 localization to stress granules. We induced oxidative stress with sodium arsenite or endoplasmic reticulum (ER) stress with thapsigargin, both of which have been previously observed to robustly recruit YTHDF2 and m^6^A-modified mRNAs to stress granules ^17,58^. Stress granules were identified using TIAR (Figure 4A). As previously reported ^17,58,59^, TIAR redistributed from a predominantly nuclear localization in unstressed cells to cytoplasmic puncta following either treatment (Figure 4D). We then quantified both the number of TIAR-positive granules and calculated the recruitment score of YTHDF2 into these compartments.

Here, the two variants diverged. Under oxidative stress, only Tyr153His significantly increased YTHDF2 recruitment to stress granules, while Tyr68His showed no significant change relative to ΔDF1/ΔDF3 (Figure 4E). This same pattern is observed under ER stress, where only Tyr153His significantly increased YTHDF2 recruitment, while Tyr68His showed no significant change (Figure 4G). The two variants also differed in their effects on stress-granule formation. Tyr153His significantly increased stress-granule number under both stress conditions, whereas Tyr68His did not alter stress-granule number under either condition (Figure 4F-4H).

Together, these data reveal distinct condensate phenotypes for the two hyperactive variants. Both Tyr68His and Tyr153His show increased partitioning of YTHDF2 into P-bodies, without altering P-body number, consistent with the enhanced degradation function we detected (Figure 2). Under stress, however, Tyr153His displays a broader effect, promoting both YTHDF2 recruitment to stress granules and granule formation itself across two mechanistically distinct stress conditions, while Tyr68His shows no significant effect on either YTHDF2 recruitment or granule number under either stress condition. These localization differences suggest that individual IDR residues can differentially regulate YTHDF2 condensate partitioning.

## DISCUSSION

In this study, we used CRISPR base-editor screening to resolve how the YTHDF2 intrinsically disordered region regulates m^6^A-modified transcripts at residue-level resolution. By densely mutagenizing full-length YTHDF2 in a sensitized ΔDF1/ΔDF3 background, we transformed the analysis of the IDR from broad fragment-level mapping into single-residue functional interrogation. This approach uncovered residues that, when mutated, promote the degradation of m^6^A-modified transcripts by disrupting YTHDF2 association with inhibitory protein partners and modulating recruitment to cytoplasmic RNA compartments. Thus, our study shows that the YTHDF2 IDR function is governed by discrete regulatory residues that tune how efficiently m^6^A recognition is converted into mRNA decay (Figure 4I).

This residue-level resolution revealed functional information that sequence-based predictors alone cannot capture. Tyr153 mapped within a predicted prion-like and droplet-promoting segment, indicating that some regulatory sites align with IDR sequence grammar (Figure 2B). By contrast, Tyr68 was among the strongest screen-selected residues but lacked prominent predicted prion-like, condensate-promoting, binding-propensity, or annotated post-translational modification features (Figure 1H, 2B). Thus, the YTHDF2 IDR contains functional residues that are only partially predictable from intrinsic sequence properties. Base-editor screening, therefore, provides an experimental platform to identify regulatory sites embedded within disordered regions that would otherwise be missed by computational annotation.

The identification of Tyr68 and Tyr153 is particularly notable because both residues are conserved across the YTHDF paralog family, suggesting that these positions may contribute to shared regulatory features of the YTHDF family (Figure S2B). Moreover, Tyr68His and Tyr153His enhanced degradation of the same CDS m^6^A-enriched transcript class that was stabilized upon YTHDF2 depletion, indicating that these mutations increase the efficiency of the YTHDF2-regulated decay program. Thus, Tyr68 and Tyr153 behave as conserved IDR regulatory nodes that act allosterically to tune YTHDF2 activity downstream of RNA recognition. This finding expands the functional architecture of YTHDF proteins beyond the YTH domain and shows that conserved residues within the disordered N-terminal region can control reader activity.

Notably, although YTHDF2 depletion and the Tyr68His/Tyr153His mutations have opposite effects on mRNA turnover, both perturbations impaired cellular fitness. This convergence suggests that YTHDF2-mediated decay must be maintained within a narrow functional range, where both insufficient and excessive degradation of YTHDF2-sensitive transcripts are detrimental.

Mechanistically, our data indicate that Tyr68 and Tyr153 inhibit YTHDF2 through association with a specific set of RNA-binding proteins, most prominently IGF2BP3. IGF2BP1, IGF2BP2, and IGF2BP3 have been described as m^6^A readers that stabilize methylated transcripts, placing them functionally opposite to YTHDF2 ^55,60–62^. Our finding that IGF2BP proteins interact with YTHDF2 suggests that these reader classes may not act only as independent or competing m^6^A-binding modules. Instead, IGF2BPs may also influence m^6^A-dependent RNA fate indirectly by regulating YTHDF2 protein interactions, localization, or access to decay-competent compartments. However, we cannot exclude that other effectors, not captured by our proximity-labeling of wild-type YTHDF2, also contribute to the hyperactive phenotype.

The localization data further support this model and link the hyperactive decay phenotype to RNA granule partitioning. P-bodies contain translationally repressed mRNAs and multiple components of the mRNA decay machinery ^63,64^, and have been proposed as cytoplasmic sites where degradation of m^6^A-marked transcripts occurs ^16,41,42^. We therefore reasoned that hyperactive YTHDF2 mutants might show increased access to these decay-associated compartments. Consistent with this idea, Tyr68His and Tyr153His increased YTHDF2 recruitment to P-bodies without increasing P-body number, indicating that these mutations alter YTHDF2 partitioning rather than globally affecting P-body assembly. Only Tyr153His significantly increased YTHDF2 recruitment to stress granules, under both oxidative and ER stress, and additionally promoted granule formation across both conditions, whereas Tyr68His showed no significant effect on either recruitment or granule number. Together, these findings suggest that Tyr68 and Tyr153 regulate a localization program that controls YTHDF2 access to cytoplasmic RNA granules, including compartments linked to mRNA repression and decay.

The convergence of Tyr68His and Tyr153His on a similar hyperactive phenotype is consistent with the multivalent nature of IDR-mediated interactions. Rather than acting through a single linear motif, IDR interaction surfaces are often distributed across multiple residues or short sequence elements, such that mutation of distinct sites can perturb binding to the same partner. Similar distributed IDR logic has been described in yeast, where multiple intrinsically disordered elements cooperate to regulate the same protein interaction ^65^. Tyr68 and Tyr153 may therefore represent separable elements of a shared YTHDF2-IGF2BP3 regulatory surface. At the same time, the screen identified additional regulatory residues. These residues may therefore correspond to other elements of the same multivalent interaction surface, as well as to distinct regulatory modules controlling YTHDF2 localization or context-dependent activity.

An important open question is whether Tyr68 and Tyr153 are dynamically regulated by post-translational modification (PTMs). PTMs within intrinsically disordered regions are thought to modulate phase separation, fine-tune interaction networks, and enable rapid, context-dependent spatial reorganization ^66^. In line with these observations, O-GlcNAcylation of YTHDF1 and YTHDF3 at IDR sites (primarily Ser196 in YTHDF1, with additional sites at Ser157, Ser197, and Ser198) is associated with stress-granule assembly and disassembly. Similarly, phosphorylation of the YTHDF1 interactor FMRP modulates the strength of the YTHDF1-FMRP interaction and its partitioning into condensates, linking a single PTM event to translational outcomes ^67^, and showing that PTM-dependent regulation of YTHDF-associated interactions and condensate behavior is an established mode of control within this protein family. Tyr68 and Tyr153 are both tyrosine residues and could, in principle, be regulated by phosphorylation. However, neither site has been annotated as phosphorylated in available proteomic datasets ^37^, largely because the surrounding IDR sequence lacks nearby tryptic cleavage sites, which may limit the detection of peptides in conventional trypsin-based phosphoproteomics ^68^. Thus, Tyr68 and Tyr153 may act directly as regulatory residues, or they may define a local interaction surface whose regulation has been missed because of limited proteomic coverage. Future studies using targeted mass spectrometry, alternative protease digestion will be needed to determine whether phosphorylation or other PTMs modulate this IDR regulatory surface.

Together, our study defines how the disordered YTHDF2 IDR, long implicated in regulating YTHDF2 activity, controls m^6^A-dependent mRNA decay at residue-level resolution. CRISPR base-editor screen identified several functional sites and validated Tyr68 and Tyr153 as conserved IDR regulatory residues that inhibit YTHDF2 activity by maintaining specific protein-protein interactions and limiting YTHDF2 partitioning into cytoplasmic RNA compartments. Disruption of this inhibitory surface releases YTHDF2 into a hyperactive state, increasing recruitment to decay-associated RNA compartments and accelerating degradation of m^6^A-modified transcripts. More broadly, our work extends base-editor screening from structured domains, disease variants, and defined regulatory sites to intrinsically disordered disordered regions. We anticipate that applying this strategy across RNA-binding proteins with intrinsically disordered regions will reveal general principles by which disordered domains encode regulatory specificity, protein-interaction logic, and context-dependent control of RNA fate.

## RESOURCE AVAILABILITY

All materials and resource requests should be directed to and will be fulfilled by the lead contact Sara Zaccara.

## DATA AND CODE AVAILABILITY

Data in this manuscript will be shared by the lead contact upon request. Data supporting these findings will be made available on public databases (GEO, Proteomexchange) upon publication or upon request.

## ACKNOWLEDGMENTS

We thank Samie Jaffrey and members of the Jaffrey lab at Weill Cornell Medicine for discussion. We also thank Chaolin Zhang, Xuebing Wu, and Anna-Lena Steckelberg groups at Columbia University for discussion. S.Z. is supported by 1DP2HD118273 and the 1K22CA258954. This research was funded in part through the HICCC-Multi-Investigator Planning Grant Award.

## AUTHOR CONTRIBUTIONS

S.Z. and B.R. conceived the project. B.R. performed most of the experiments and analyses. S.Z. supervised the project. S.Z. and J.G. performed and supervised the μMAP analyses. D.N. performed protein binding assays. D.S. assisted with library preparation and protein interaction analysis. R.Z. assisted with validation, supervised by B.R. S.Z. and B.R. wrote the manuscript with input from all authors.

## SUPPLEMENTARY INFORMATION

**Table S1:** Base-editor screen results for ABE and CBE screens (sgRNA fold change, Z scores, and predicted mutations). Related to Figure 1.

**Table S2:** TimeLapse-seq gene-level k_deg values, m^6^A stoichiometry, and metagene region assignments for ΔDF1/ΔDF3, Tyr68His, Tyr153His, and ΔDF1/ΔDF3+shDF2. Related to Figure 2.

**Table S3:** Endogenous YTHDF2 μMAP proximity-labeling proteomics, differential enrichment over IgG in paraformaldehyde- and methanol-fixed conditions. Related to Figure 3.

**Table S4:** Oligonucleotides and sgRNA sequences used in this study.

## MATERIALS AND METHODS

### Cell culture

HeLa and HEK293T cells (ATCC) were maintained in Dulbecco’s Modified Eagle Medium (DMEM; Fisher Scientific, MT10090CV) supplemented with 10% fetal bovine serum (FBS), 100 U mL⁻¹ penicillin, and 100 μg mL⁻¹ streptomycin at 37°C in a humidified incubator with 5% CO₂. Cells were passaged using TrypLE Express (Gibco, Thermo Fisher, 12604021) according to the manufacturer’s instructions.

### Generation of ΔYTHDF1/ΔYTHDF3 cells

YTHDF1/YTHDF3 double-knockout HeLa cells were generated by sequential CRISPR-Cas9-mediated genome editing using Cas9 ribonucleoprotein (RNP) delivery to avoid stable Cas9 expression. In vitro-synthesized gRNAs targeting YTHDF3 (Table S4; Integrated DNA Technologies) were complexed with recombinant Alt-R S.p. Cas9-GFP V3 (10008100; Integrated DNA Technologies) according to the manufacturer’s instructions and nucleofected into HeLa cells using the SE Cell Line 4D-Nucleofector Kit and program CN-114 (Lonza). Three days after nucleofection, GFP-positive cells were isolated by fluorescence-activated cell sorting and plated as single cells. Individual clones were expanded and screened for YTHDF3 loss by immunoblotting. A validated YTHDF3-knockout clone was then subjected to the same procedure using gRNAs targeting YTHDF1. Individual clones were expanded and screened for loss of both YTHDF1 and YTHDF3 by immunoblotting. A validated YTHDF1/YTHDF3 double-knockout (ΔYTHDF1/ΔYTHDF3) clone was used for all subsequent experiments.

### Generation of Doxycycline-inducible shYTHDF2 cells

To generate ΔYTHDF1/ΔYTHDF3 cells with inducible YTHDF2 depletion, a doxycycline-inducible lentiviral vector (Addgene, 21915) encoding a short hairpin RNA (shRNA) targeting *YTHDF2* (Table S4) was introduced by lentiviral transduction (see Lentivirus production). Transduced cells were selected with 1μg/mL puromycin for 3 days to establish stable cell populations. YTHDF2 knockdown was induced by doxycycline (2 μg/mL) for 48 hours before downstream experiments, and depletion efficiency was confirmed by immunoblotting.

### Lentivirus production and transduction

Lentiviruses were produced in HEK293T cells by transient transfection of the plasmid of interest together with the packaging plasmid psPAX2 (Addgene 12260) and the envelope plasmid pMD2.G(Addgene 187440) using Lipofectamine (LipoD293; Signagen, SL100668) according to the manufacturer’s instructions. Briefly, HEK293T cells were transfected in 10-cm culture dishes, viral supernatants were collected at 48 and 72 h post-transfection, clarified by centrifugation at 300 × g for 5 min and filtered through a 0.45-μm syringe filter. Lentiviral particles were concentrated using PEG-it Virus Precipitation Solution (Fisher Scientific, LV825A1) according to the manufacturer’s instructions, resuspended in Opti-MEM (Thermo Fisher Scientific, 31985070), aliquoted, and stored at −80°C until use. The concentration of lentiviral particles was estimated using the Lenti-X GoStix Plus assay (Takara, 631280) according to the manufacturer’s instructions. Briefly, 20 μL of lentiviral supernatant was added to a GoStix cassette, followed by the supplied Chase Buffer. After incubation for 10 min at room temperature, the cassette was scanned using the Lenti-X GoStix smartphone app., and lentiviral p24 abundance was reported as a GoStix Value. Infectious titers (infectious units per milliliter, IFU/mL) were estimated from the measured p24 concentration using the manufacturer’s conversion algorithm and were used to normalize viral input across experiments.

For lentiviral transduction, target cells were seeded at 5 × 10⁵ cells per well in 12-well plates and incubated with the lentivirus of interest in a final volume of 1mL complete culture media. 1ug/mL of polybrene was added to help transduction. Plates were centrifuged at 640 × g for 90 min at 16C temperature. At 48h after transduction, cells were transferred to a 6 well plates and culture in complete media supplemented by 1 μg/mL puromycin (Gemini Bio Products, Fisher Scientific, 507533041) until all cells in the untrasduced control population had died. Surviving cells were expanded as stable clonal cell populations, and mutation or protein depletion was confirmed by Sanger sequencing or Western blot, respectively.

### Transient siRNA-mediated knockdown

Transient gene knockdown was performed using small interfering RNAs (siRNAs; Table S4) as also described in ^11^. 250.000 ΔYTHDF1/ΔYTHDF3 cells were seeded 24 hours prior to transfection in a 6 well. 3uL of 10uM siRNA together with 3.6uL PepMute Transfection Reagent (Signagen, 504845) were added to 100uL PepMute Transfection Buffer. Complexes were incubated for 10 min at room temperature before being added to the cells. A second round of transfection was performed after 48 hours, and cells were harvested on day 4. Knockdown efficiency was confirmed by immunoblotting.

### WST-1 proliferation assay

Cell proliferation was assessed using the WST-1 Cell Proliferation Reagent (Sigma-Aldrich, 05015944001). Cells were seeded in 96-well plates. At the indicated time points, 10 μL WST-1 reagent was added directly to each well and incubated for 30 minutes at 37°C according to the manufacturer’s instructions. Absorbance was measured at 450 nm using a microplate reader. Background absorbance from media-only wells was subtracted prior to analysis.

### Generation of clonal populations

Clonal YTHDF2 edited cell lines were generated in the ΔYTHDF1/ΔYTHDF3 background by lentiviral transduction of a plasmid encoding either CBE or ABE base editor and the appropriate Tyr68His and Tyr153His sgRNAs, respectively. Because the parental ΔDF1/ΔDF3 cell line was generated by transient Cas9 RNP delivery, these cells did not constitutively express Cas9 before introduction of the lentiviral base-editor construct. Following puromycin selection, as described above, cells were plated at limiting dilution into 96-well plates to obtain single-cell-derived colonies. Individual clones were expanded and screened for the editing by Sanger sequencing as follow. Genomic DNA was extracted using QuickExtract (VWR, QE09050) according to the manufacturer’s instructions. Genomic regions spanning the edited loci were PCR-amplified using primers flanking the target sites (Table S4). PCR products were purified and sent for Sanger Sequencing (Genewiz). Sequencing chromatograms were aligned to the wild-type YTHDF2 reference sequence using Snapgene to confirm successful editing. Clones with the expected mutations were used for subsequent experiments. In some cases, the used gRNA caused additional editing upstream or downstream of the preferred editing site. Clones were excluded from the subsequent experiments when the additional editing caused a change in the aminoacidic sequence.

### Base editor screen

#### Library design and construction

A YTHDF2 base-editing library was designed and constructed at the Genetic Perturbation Platform of the Broad Institute of MIT and Harvard following the guidelines previously reported ^30^. Briefly, the principal Ensembl transcript of YTHDF2 (ENST00000373812.8) was selected to obtain the transcript and protein sequences and used to annotate each sgRNA with its predicted edits. We included all sgRNAs targeting the coding sequence in both orientations; we also included all sgRNAs for which the start was up to 30 nucleotides into the intron and UTRs. The four nucleotides immediately downstream of each 20-nt protospacer were retained as the PAM sequence. Guides containing BsmBI recognition sites or poly-T stretches (TTTT) were excluded. Promiscuous guides were also removed, defined as guides whose PAM-proximal 18-nt sequence had five or more genomic off-target matches with up to one mismatch. The final library contained 816 YTHDF2-targeting sgRNAs, together with 192 positive-control and 1,000 negative-control sgRNAs. Oligonucleotide pools were synthesized by CustomArray, with BsmBI sites and cloning overhangs appended to each protospacer for insertion into the sgRNA-expression vector. The used vectors are pRDA_429, NG-ABE8e (Addgene #179098); pRDA_336, NG-BE3 (Addgene #179095).

#### Pooled base-editing screens and sequencing

Cells were transduced with the pooled lentiviral sgRNA library in two biological replicates. Lentiviral titers were determined by transducing cells with serial virus volumes, followed by paired puromycin-selected and unselected cultures. A viral dose yielding 30-50% transduction efficiency, corresponding to an MOI of approximately 0.35-0.70, was used for screening.

Because the titer of all-in-one base editor viruses was low, cells were plated in polybrene-containing media with 1.5 × 106 cells per well in a 12-well plate. Plates were centrifuged for 2 h at 640 x g, after which 2 mL of media was added to each well. Plates were then transferred to an incubator for 4-6 h, after which virus-containing media was removed, and cells were pooled into flasks. Puromycin was added 2 days post-transduction and maintained for 5-7 days to ensure complete removal of non-transduced cells. Upon puromycin removal, cells were split at a representation of at least 1000 cells per sgRNA and passaged every 2-4 days for an additional 21 days to allow sgRNAs to enrich or deplete; cell counts were taken at each passage to monitor growth.

Genomic DNA was isolated at the indicated time points. sgRNA cassettes were amplified from genomic DNA using barcoded PCR primers, with no more than 10 μg genomic DNA per 100-μL PCR reaction. Amplified libraries were purified with AMPure XP beads and sequenced on an Illumina HiSeq 2500 High Output platform with a 5% PhiX spike-in at the Genetic Perturbation Platform of the Broad Institute of MIT and Harvard facility.

### Bacterial expression and purification of YTHDF2 and its mutants

YTHDF2, YTHDF2 Trp432Ala, YTHDF2 Tyr68His, and YTHDF2 Tyr153His were cloned in the pTXB1 expression plasmid (NEB # N6707S) using Gibson Cloning. Proteins were then purified using the IMPACT system (New England Biolabs E6901S) according to the manufacturer’s guidelines, with minor adjustments made to the original protocol. For protein production, plasmids constructs expressing either YTHDF2 or its mutants were introduced into *E.coli* strain C2566H. Overnight cultures were diluted to an initial OD600 nm of 0.1 in 200 mL of LB medium and incubated at 37°C with shaking until reaching an OD600 nm of about 0.5. Cultures were then cooled on ice and subsequently induced with 0.1 mM Isopropyl β-D-1-thiogalactopyranoside (IPTG), followed by incubation at 30°C for 6 h with shaking. Cells were collected by centrifugation. Cell pellets were resuspended in lysis buffer (20 mM HEPES, 800 mM NaCl, 1 mM EDTA, 0.5% Triton-X, 0.5 mM Lysozyme, 100X Protease Inhibitor), incubated on ice for 15 minutes, and followed by disruption by sonication for 6 minutes, 30-second pulses, and 20% amplitude. After centrifugation to clarify the lysates, the supernatants were loaded onto Poly-Prep Chromatography Columns (Bio-Rad #7311550) containing 1 mL of chitin resin (New England Biolabs S6651S) pre-equilibrated with 10 mL of washing buffer (20 mM HEPES, 800 mM NaCl, 1 mM EDTA, 0.5% Triton-X) and left to incubate for 20 minutes. The columns were washed with 10 mL of wash buffer before initiating the on-column cleavage. To release the proteins, the resin was incubated at 4°C with cleavage buffer (wash buffer supplemented with 50 mM DTT) overnight. The proteins were then eluted with 1 mL of wash buffer, and protein enrichment and isolation were verified by SDS-PAGE and Coomassie staining. Eluted proteins were loaded onto Pierce Protein Concentrators with a 50 kDa cutoff (Thermo Scientific 88504) for buffer exchange into the final working and storage buffer (20 mM HEPES, 150 mM NaCl, 20% Glycerol). The proteins were quantified using an Albumin standard curve and aliquots were stored at -80°C.

### Electrophoretic mobility shift assay (EMSA)

The binding of YTHDF2 and its mutants to RNA was evaluated by fluorescence electrophoretic mobility shift assay, with considerations adapted from ^69^. Binding reactions were prepared in a 10 ul volume containing 50 nM of a methylated fluorescent RNA (Table S4), 5X Binding Buffer (50 mM HEPES, 125 mM KCl, 0.5 mM EDTA, 50% Glycerol, 0.25% Triton-X), 0.5 uM of the same RNA but unmethylated, as competitor, 20 U of RNase inhibitor (Invitrogen AM2696), and increasing amounts of purified YTHDF2 or mutant protein. The mixtures were incubated at room temperature for 30 minutes in the dark. Samples were then loaded into a cast native 6% acrylamide TBE gel after adding Orange G loading dye (Fisher Scientific AAJ60562AC). The gel was pre-run for 30 minutes at 80 V and then run for approximately 30 minutes at the same voltage with cold 0.5X TBE buffer and in the dark. Gels were then incubated for 3 minutes with Sybr Gold (Thermo Fisher S11494) and imaged directly using a Typhoon scanner with the Cy2 fluorescence setting.

### Antibody-directed μMAP proximity labeling

Antibody-directed μMAP proximity labeling was performed as previously described by Geri and colleagues ^48^, with the following modifications. For each replicate, four 10 cm dishes of HeLa cells were fixed with either 4% paraformaldehyde or ice-cold methanol for 15 min at 4°C with gentle agitation in the cold room. Cells were washed twice with 1× PBS and blocked for 30 min in 1× PBS supplemented with 0.2% Triton X-100 and 2% heat-inactivated FBS. Cells were then incubated overnight with a YTHDF2-specific primary antibody (Proteintech; 24744-1-AP) diluted in 5 mL blocking buffer per 10 cm dish. As a negative control, four plates were incubated in parallel with normal rabbit IgG (Cell Signaling Technology, 2729S). The following day, cells were washed three times with 1× PBS containing 0.2% Triton X-100. A photocatalyst-conjugated goat anti-rabbit secondary antibody was prepared as previously described and incubated with cells for 1 h in the dark at a concentration fourfold higher than that used for the primary antibody. Cells were then washed once with 1× PBS containing 0.2% Triton X-100, followed by two washes with 1× PBS. All washes were performed in the dark. Cells were incubated with biotin-labeling reagent in 1× PBS for 10 min in the dark and then irradiated with blue light for 5 min to induce proximity-dependent labeling. After labeling, cells were lysed directly on the plate by adding 1 ml RIPA buffer supplemented with 1% SDS per 10 cm dish and heating at 90°C for 5 min. Cells were scraped, transferred to 1.5 ml tubes, and sonicated at 90% amplitude for 1 min. Lysates were clarified by centrifugation at 10,000 × g for 5 min. Streptavidin enrichment of biotinylated proteins was performed as previously described. Enriched proteins were analyzed by mass spectrometry using a Timstof Pro2.

### Endogenous IGF2BP3 co-immunoprecipitation

For endogenous co-immunoprecipitation experiments, 5 × 10⁶ cells were seeded in 15-cm dishes 24 h prior to lysis. Cells were washed with ice-cold PBS and lysed on ice in CHAPS lysis buffer (1% CHAPS, 25 mM Tris-HCl, pH7.5, 20 mM NaCl, 2 mM MgCl₂, 0.5% glycerol, and 0.02% SDS) supplemented with protease inhibitors (Roche, 11836153001), phosphatase inhibitors (Roche, 4906845001), and RNase inhibitor (Thermo Scientific, AM2696). Lysates were incubated for 15 min at 4°C with rotation and clarified by centrifugation at 10,000 × g for 5 min. Protein concentrations were determined using the Pierce BCA Protein Assay Kit (Thermo Fisher Scientific, 23225), and 10% of each lysate was reserved as input. Equal amounts of protein were incubated with 2 μg antibody (IGF2BP3, Proteintech,14642-1-AP) per mg lysate for 2 h at 4°C with rotation before addition of Protein A/G magnetic beads (Pierce, PI88802). Immune complexes were captured for 30 min at 4°C, washed once with high-salt CHAPS buffer (150 mM NaCl) followed by three washes with standard CHAPS buffer (20 mM NaCl), and eluted by boiling for 5 min in 4× Laemmli sample buffer with DTT. Eluted proteins were resolved by SDS-PAGE and analyzed by immunoblotting.

### Proximity labeling using YTHDF2-TurboID fusion constructs

Plasmids encoding YTHDF2-TurboID and YTHDF2 mutant variants (W432A, Y68H, and Y153H) were transformed into competent bacteria, plated, and grown overnight. Single colonies were inoculated into cultures for plasmid preparation. Hela ΔYTHDF1/3 cells were transfected with 12 μg of plasmid DNA using 23 μL of FuGENE HD transfection reagent (Promega, E2311) diluted in 1.15 mL of incomplete DMEM per 100mm dish.Twenty-four hours post-transfection, biotinylation was initiated by adding fresh DMEM containing 50 μM of biotin (Sigma-Aldrich, B4501) per dish, followed by incubation at 37°C for 10 minutes. The reaction was terminated by removing the media, transferring dishes to ice, and washing the cells five times with 1X ice-cold PBS.Cells were scraped and lysed with 500 µl of RIPA buffer containing protease inhibitors and incubated on ice for 15 minutes. Cells were spun down and the supernatant was collected for BCA assay protein quantification. Following quantification, an input fraction of ∼30μg of total protein was reserved and the remaining lysate was used for streptavidin pulldowns. Biotinylated proteins were enriched by adding an equal amount of RIPA buffer-exchanged streptavidin-conjugated magnetic beads (Thermo Fisher Scientific, 88817, 25µl/300µg of protein) per sample. Samples were incubated for 1 hour at 4°C with rotation. Beads were washed once with RIPA buffer supplemented with 300 mM NaCl, followed by two washes with standard RIPA buffer. Biotinylated proteins were eluted by boiling beads in 40 μL of 1.5× protein loading buffer supplemented with 2 mM biotin and 20 mM DTT at 95°C for 10 minutes. Beads were pelleted using a magnetic rack and eluates were collected for western blot.

### Immunofluorescence

Cells were seeded onto Lab-Tek chambered coverglass slides (Thermo Fisher, 171080) and fixed with 4% paraformaldehyde in PBS for 15 minutes at room temperature. Following fixation, cells were permeabilized with 0.02% Triton X-100 in PBS for 20 minutes and blocked with 2% FBS in PBS for 30 minutes. Samples were incubated overnight at 4°C with primary antibodies diluted in blocking buffer at the following concentrations: anti-YTHDF2 (Proteintech, 24744-1-AP) 1:100; anti-EDC4 (Santacruz, sc-374211) 1:50; and anti-TIAR (Cell Signaling, 5137) 1:50. Slides were washed three times with PBS, and incubated with species-specific Alexa Fluor-conjugated secondary antibodies (anti-mouse A10037; anti-rabbit A32790) diluted 1:1000 in blocking buffer for 1 h at room temperature. The slides were protected from light during from secondary antibody addition onwards. Nuclei were stained using Hoechst (Thermo Scientific, 62249) diluted 1:10000 in PBS for 10 minutes at room temperature. Samples were mounted using ProLong Diamond Antifade Mountant (Fisher Scientific, P36965) mounting medium. Images were acquired using a Nikon NIS-Elements Eclipes Ti2 confocal microscope. Quantitative image analysis was performed using the General Analysis 3 (GA3) module in Nikon NIS-Elements software using identical acquisition settings for all samples within an experiment.

### P-bodies staining and quantification

P-bodies were detected by immunofluorescence staining for the P-body marker EDC4. Cells were co-stained with anti-EDC4 (1:50) (Santacruz, sc-374211) and anti-YTHDF2 (1:100) (Proteintech, 24744-1-AP) antibodies as described above. EDC4-positive puncta were identified using the GA3 module in NIS-Elements based on object intensity and size thresholds. P-body number per cell and YTHDF2 fluorescence intensity within EDC4-positive objects were quantified for individual cells. YTHDF2 enrichment in P-bodies was calculated as the mean YTHDF2 fluorescence intensity within EDC4-positive puncta relative to the mean fluorescence intensity in the corresponding cell. Identical segmentation and detection thresholds were applied to all conditions within each experiment.

### Stress granule induction and staining

To induce oxidative stress, cells were treated with 0.5 mM sodium arsenite for 30 min at 37°C. To induce ER stress, cells were treated with 1 μM thapsigargin for 1 h at 37°C. Following treatment, cells were fixed and processed for immunofluorescence as described above.

Stress granules were detected using an anti-TIAR antibody (1:50), together with anti-YTHDF2 (1:100) (Proteintech, 24744-1-AP). TIAR-positive puncta were identified using the GA3 module in NIS-Elements based on intensity and size thresholds. Stress-granule number, and fluorescence intensity were quantified per each individual cell. YTHDF2 recruitment at stress granules was calculated as the mean YTHDF2 fluorescence intensity within TIAR-positive puncta relative to the mean fluorescence intensity in the corresponding cell. Identical segmentation and detection thresholds were applied to all conditions within each experiment.

### Western blot

Cells were washed once with PBS and lysed in RIPA buffer (50 mM Tris-HCl, pH 7.5, 150 mM NaCl, 1% NP-40, 0.5% sodium deoxycholate, and 0.1% SDS) supplemented with 1× protease inhibitor cocktail (Roche, 11836153001). Lysates were sonicated (20% amplitude, 5 s on/5 s off for a total sonication time of 15 s) and clarified by centrifugation at 10000 × g for 5 min at 4°C. Protein concentration was determined using the Pierce BCA Protein Assay Kit (Thermo Fisher Scientific, PI88802), and equal amounts of protein (15 μg) were mixed with 4× Laemmli sample buffer containing 1mM DTT and denatured at 95°C for 5 min. Proteins were resolved by SDS-PAGE on 4-12% Bis-Tris gels and transferred to PVDF membranes for 1 hour by wet transfer.

Membranes were blocked in 5% milk in TBS-Tween for 30 minutes at room temperature and probed with primary antibodies overnight at 4°C at the following dilutions anti-YTHDF2 (Proteintech, 24744-1-AP) 1:3000; anti-IGF2BP3 (Cell Signaling Technology, 57145S) 1:1000; anti GAPDH (Proteintech, 10494-1-AP) 1:5000. Membranes were washed three times with TBST for 5 minutes each and incubated with HRP-conjugated secondary antibodies (Thermo Fisher Goat anti-Mouse IgG (H+L) Secondary Antibody, HRP, 31430 or Thermo Fisher Goat anti-Rabbit IgG (H+L) Secondary Antibody, HRP, 31460) diluted 1:5000 in 5% milk for 1 hour at room temperature. Membranes were washed three times with TBST for 5 minutes each. Bands were visualized by enhanced chemiluminescence using Immobilon Western Chemiluminescent HRP Substrate (Millipore-Sigma, WBKLS0050) or SuperSignal™ West Atto Ultimate Sensitivity Substrate (Thermo Fisher, A38555) depending on signal intensity and imaged on a BioRad ChemiDoc system.

### Timelapse-Seq

#### 4sU labeling and RNA isolation

Cells were seeded and subjected to siRNA-mediated depletion, where indicated, as described above. Newly synthesized RNA was labeled by the addition of 4-thiouridine (4sU; Fisher Scientific, AAJ60679MD) to the culture medium at a final concentration of 500 μM for 2 h. Cells were washed with PBS and lysed directly in TRIzol reagent (Thermo Fisher Scientific, 15596018). Total RNA was isolated according to the manufacturer’s instructions and resuspended in nuclease-free water. RNA concentration was determined using the Nanodrop.

#### Timelapse chemical conversion

TimeLapse conversion of 4sU-labeled RNA was performed as previously indicated ^70^. Briefly, for each sample, 2 μg total RNA was brought to a final volume of 8.7 μL in DEPC-treated water. A conversion master mix containing 0.84 μL 3 M sodium acetate, pH 5.2, 0.2 μL 0.5 M EDTA, pH 8.0, 1.3 μL TFEA, and 12.7 μL DEPC-treated water was added to each sample. Reactions were initiated by addition of 1.3 μL freshly prepared 192 mM sodium periodate, mixed, and incubated for 1 h at 45°C in a thermocycler with a heated lid. Converted RNA was purified by addition of an equal volume of RNAClean XP beads (Beckman Coulter, A63987), followed by incubation for 10 min at room temperature. Beads were captured on a magnetic rack, washed twice with freshly prepared 80% ethanol, air-dried briefly, and eluted in 18 μL DEPC-treated water. Converted RNA was reduced by addition of 2 μL 10× reducing buffer containing 100 mM Tris-HCl, pH 7.4, 100 mM DTT, 1 M NaCl, and 10 mM EDTA, followed by incubation for 30 min at 37°C. RNA was purified again using RNAClean XP beads and eluted in 12 μL DEPC-treated water as above mentioned. RNA concentration and integrity were assessed after conversion using an Agilent Bioanalyzer and the RNA 6000 Pico Kit (Agilent Technologies, 5067-1513). Samples showing substantial RNA degradation or insufficient yield were excluded from library preparation.

#### Library preparation

RNA libraries were prepared from 1 μg of non-converted RNA and 2 μg of converted RNA. ERCC RNA spike-in controls (Thermo Fisher, 4456740) were added to non-converted samples at a 1:100 dilution (2 μL per 1 μg of total RNA). Ribosomal RNA was depleted using the NEBNext rRNA Depletion Kit (New England Biolabs, E7400), and libraries were prepared using NEBNext Ultra™ II Directional RNA Library Prep Kit for Illumina (New England Biolabs, E7760) with the following modifications: RNA fragmentation was performed for 11 minutes for non-converted samples and 8 minutes for converted samples, and adapters were used at a 5X dilution.

### QUANTIFICATION AND STATISTICAL ANALYSIS

#### Base editor screen

##### Guide-level analysis and predicted coding consequences

Sequencing counts were analyzed separately for the AM14/NG-ABE8e and AM23/NG-BE3 screens using MAGeCK to generate guide-level LFC values using the default settings. Predicted nucleotide and amino-acid consequences were classified as nonsense, missense, or silent mutations with a defined amino-acid position. When a guide had multiple predicted coding edits, only the first valid coding mutation was used for guide-level residue assignment.

For each screen, robust z-scores were calculated from MAGeCK LFCs relative to the negative-control guide distribution:

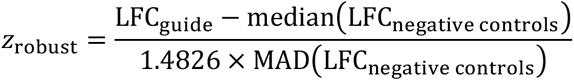

Guides at or below the 2nd percentile or at or above the 98th percentile of the screen-specific robust-z distribution were classified as top-scoring guides. Silent first-coding mutations were excluded from graphical hit displays and hit summaries because their phenotypic effects are difficult to attribute directly to altered protein sequence and require more careful follow-up evaluation.

#### TimeLapse analysis

TimeLapse-seq data were processed using the fastq2EZbakR and EZbakR suite according to the recommended pipeline ^40^. Sequencing reads were processed with the fastq2EZbakR Snakemake pipeline to quantify nucleotide-conversion events and assign reads to annotated genomic features, generating counts-binomial (cB) files for downstream analysis. Kinetic analyses were subsequently performed in R using EZbakR and bakR.

For each comparison, converted samples and matched non-converted controls were analyzed together. EZbakR objects were generated from the corresponding cB files using EZbakRData. Metabolic labeling fractions were estimated using EstimateFractions with hierarchical modeling, genotype as the grouping factor, and non-converted samples used to estimate background mutation rates. Estimated nucleotide-conversion rates for labeled and non-converted samples are reported in Fig. S2C. RNA kinetic parameters were inferred using EstimateKinetics, followed by dropout correction using CorrectDropout with the bakR strategy. Gene-level kinetic estimates were averaged and regularized across biological replicates using AverageAndRegularize. For the Tyr153Hist comparison, which included samples from two sequencing batches, both genotype and batch were included in the model (∼ genotype + batch); the other comparisons were performed within a single sequencing batch and were modeled using genotype alone (∼ genotype). Differences in RNA degradation (log_kdeg) rates between conditions were determined using CompareParameters, with genotype specified as the design factor. Quality-control, MA, and volcano plots were generated using the corresponding EZbakR functions.

Cumulative distributions of log(k_deg) were compared between genotypes using two-sided Kolmogorov-Smirnov tests, and fold-changes in per-gene k_deg were assessed using two-sided Wilcoxon signed-rank tests against a null of zero. Multiple comparisons within a given analysis were corrected using Benjamini-Hochberg (BH). Significance is denoted throughout the figures as *p < 0.05, **p < 0.01, ***p < 0.001, ****p < 0.0001; ns, not significant (p ≥ 0.05), with the specific test, correction method, and sample size (n, defined as biological replicates unless otherwise specified) stated in each figure legend.

#### μMAP analysis

##### Calculation of log2FC

Protein-level differential enrichment analysis was performed in R from DIA protein-group intensity matrices using the *MSnbase*, *DEP*, and *limma* packages. Protein groups were first converted into unique identifiers based on the reported gene and protein-group annotations, and zero-intensity values were recoded as missing values. For DIA datasets, peptide support per protein group was derived from the precursor-level matrix by collapsing unique stripped peptides and counting the number assigned to each protein group. Only proteins supported by more than two peptides were retained for downstream analysis. To accommodate the immunoprecipitation design, the two experimental arms (*low_a* and *high_a*; four replicates each in this analysis) were processed separately rather than normalized together. For each protein, the fraction of missing values was calculated independently in each arm, and proteins were retained if the proportion of missing values was below 30% in at least one arm. Proteins were also flagged according to whether they required no imputation, partial imputation, or full imputation in one arm. Within each arm, intensities were normalized by variance stabilizing normalization (*vsn*), after which missing values were imputed using the *MinProb* method. The separately normalized and imputed arm-specific matrices were then recombined into a single expression matrix for statistical testing. Differential enrichment between *low_a* and *high_a* was estimated using *limma* with an empirical Bayes moderated linear model. The moderated *logFC* values from this model were used as the protein-level log2FC measurements for downstream ranking and visualization, and P values were adjusted in the limma workflow using the Benjamini-Hochberg test.

##### Comparison with NMF

For NMF-based annotation of YTHDF2 interactors, we selected the top 25 proteins from the overlap between the methanol- and formaldehyde-based YTHDF2 pulldown datasets, restricting the analysis to shared significant interactors with positive enrichment and ranking proteins by mean log2 fold change across the two conditions. These proteins were matched to the previously published non-negative matrix factorization reference table Table1 of Youn et al., 2018 ^51^, from which the 14 NMF rank scores were extracted for each protein. To assess similarity to the YTHDF family, we computed the cosine similarity between each interactor’s 14-rank NMF profile and the profile derived from YTHDF2. Because rank 8 corresponded to the CCR4-NOT complex/P-body module, we interpreted this rank as a degradation-associated signature. The resulting data were visualized as heatmaps displaying proteins grouped by predominant NMF rank and ordered by rank 8 loading, with YTHDF-profile similarity used as a secondary ordering criterion.

## DECLARATION OF GENERATIVE AI and AI-ASSISTANT TECHNOLOGIES

During the preparation of this manuscript, the authors used ChatGPT to enhance the readability and clarity of certain sections. All content generated with the assistance of this tool was reviewed and edited by the authors, who take full responsibility for the final version of the manuscript.

---

**Figure S1:**
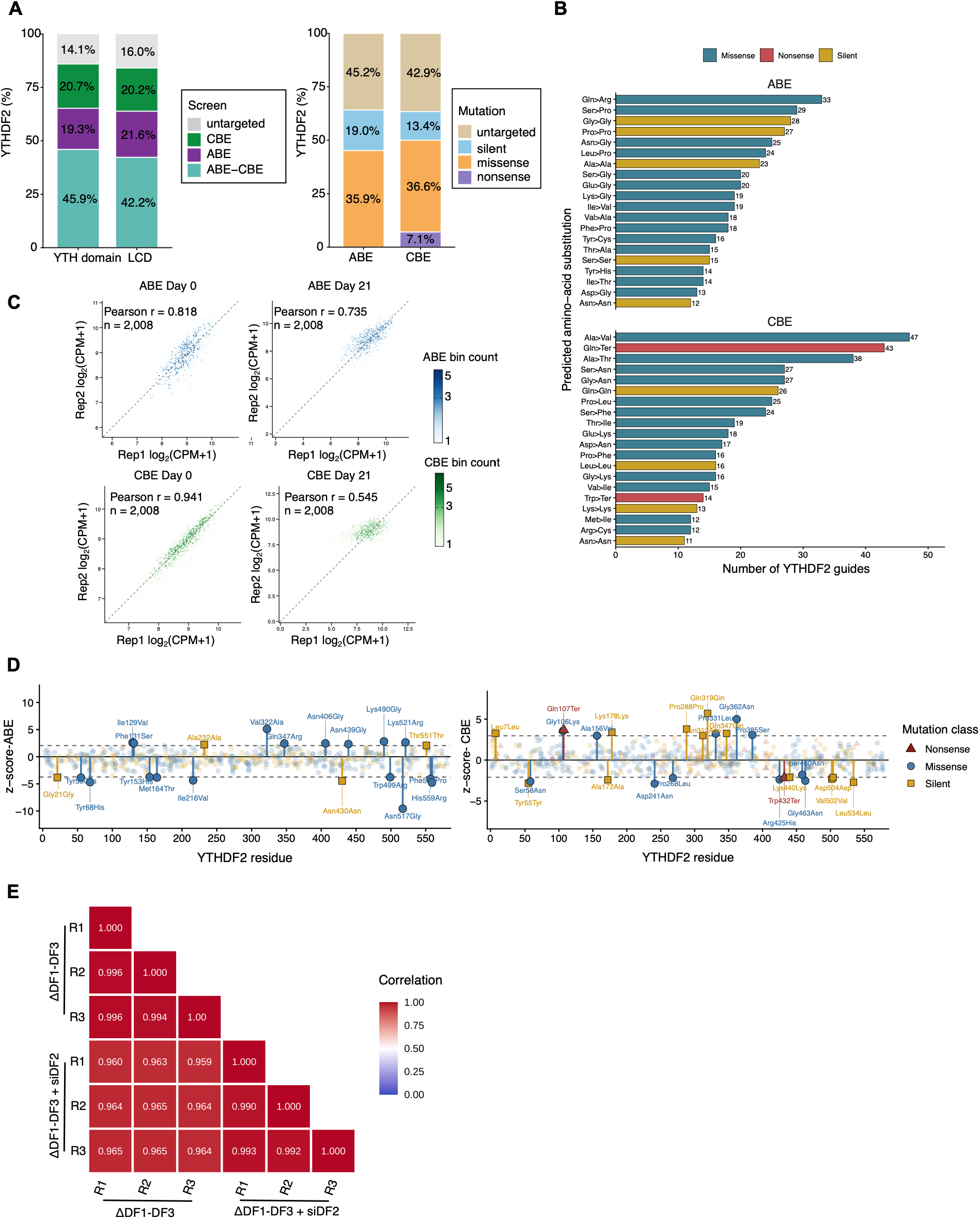
Design, coverage, and quality control of the YTHDF2 base editor sgRNA libraries. **(A)** Coverage of the YTH domain and intrinsically disordered region by ABE, CBE, or both (ABE-CBE) sgRNAs, versus untargeted residues (left); breakdown of predicted mutation type (silent, missense, nonsense) for ABE and CBE-targeted residues within YTHDF2 (right). **(B)** Number of YTHDF2-targeting sgRNAs by predicted amino-acid substitution and mutation class (missense, nonsense, silent) for the ABE (top) and CBE (bottom) libraries. **(C)** Reproducibility of sgRNA abundance in the ABE and CBE screens. Scatter plots show guide-level abundance, expressed as log2(CPM + 1), between biological replicates at day 0 and day 21 for the ABE (top) and CBE (bottom) screens. Points are colored by guide-count density. Pearson correlation coefficients are indicated for each comparison. The number of gRNAs is indicated. **(D)** Guide-level robust z-scores plotted across YTHDF2 residue position for the ABE (left) and CBE (right) screens, colored by predicted mutation class (nonsense, missense, silent). Related to Figure 1H. **(E)** Reproducibility of RNA-seq libraries. Pearson correlation matrices of log-transformed gene-level read counts, shown as log1p (counts), across RNA-seq replicates of ΔDF1/ΔDF3 and ΔDF1-DF3+siDF2 cells (62000 genes). Heatmap color indicates the Pearson correlation coefficient.

**Figure S2:**
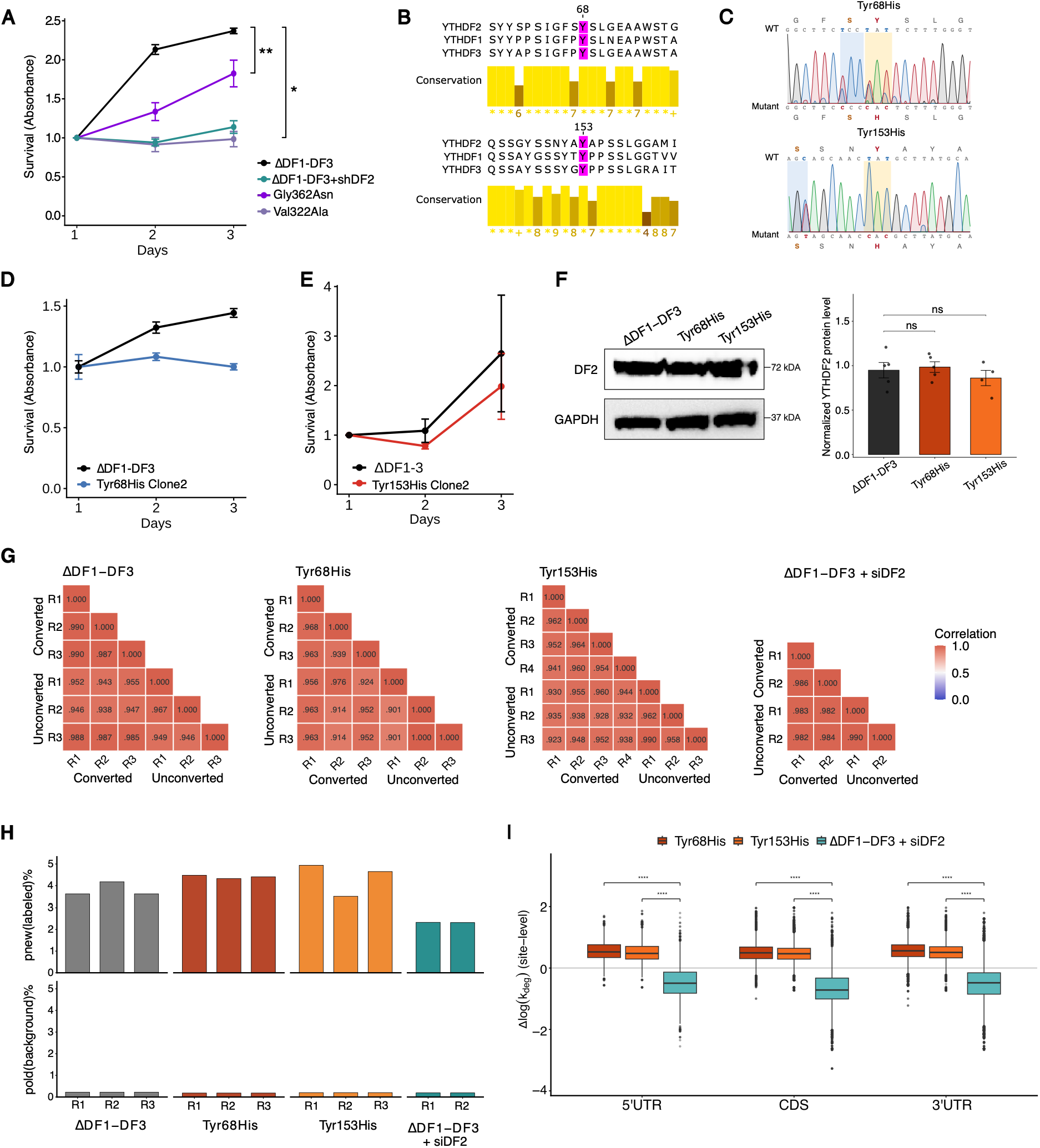
Validation and quality control for YTHDF2 IDR mutants. **(A)** Proliferation of ΔDF1/ΔDF3, ΔDF1-3+shYTHDF2 cells and cells expressing individual base-editor sgRNAs targeting candidate IDR residues predicted from the base editor screens to enhance proliferation (Gly362Asn, Val322Ala), measured by WST-1 proliferation assay over 3 days. Data are mean ± SEM of n=3 biological replicates. Neither candidate conferred a reproducible proliferation advantage under these short-term validation conditions when tested individually. (**B**) Sequence conservation of Tyr68 and Tyr153 across the YTHDF paralog family. Amino acid alignment of YTHDF1, YTHDF2, and YTHDF3 spanning the regions surrounding Tyr68 and Tyr153. Bar plots below each alignment show per-position Jalview conservation scores across the three paralogs (yellow). Bar height reflects the degree of physicochemical conservation at that alignment column, scored on Jalview’s 0-11 scale (calculated via the AMAS method), where 11 indicates complete conservation of amino acid physicochemical properties across all three sequences and lower values indicate divergence. Both residues, and their surrounding sequence context, are conserved across YTHDF1-3, suggesting these positions may represent a conserved regulatory feature of the YTHDF intrinsically disordered region. **(C)** Representative Sanger sequencing traces confirming successful base editing at Tyr68 and Tyr153 in ΔDF1/ΔDF3 cell lines. The wild-type (WT) amino acid sequence is shown for reference. The expected TAT-to-CAT codon change, corresponding to Tyr-to-His substitution, is indicated relative to the unedited wild-type sequence. The Tyr68 targeting sgRNA also produced a lower-frequency bystander edit at the adjacent serine codon (Ser67Pro) (in blue), and the Tyr153 targeting sgRNA produced a silent bystander edit at Ser150. Both upstream edits are silent; thus, they are not expected to affect the observed findings related to the missense mutation. (**D-E**) Proliferation of independently isolated Tyr68His (Clone 2) and Tyr153His (Clone2) clones relative to ΔDF1/ΔDF3 control, measured by WST-1 (Tyr68His) or Cell titer (Tyr153His) proliferation assay over 3 days. Both clones showed reduced proliferation relative to control, confirming the growth defect as a reproducible consequence of each IDR substitution rather than a clonal artifact. n=1 biological replicate. **(F)** YTHDF2 protein levels do not change in edited IDR mutant cell lines. Representative immunoblot of total YTHDF2 in ΔDF1/ΔDF3 cells expressing wild-type YTHDF2 and in Tyr68His and Tyr153His edited cell lines. GAPDH was used as a loading control. Quantification shows normalized protein levels for n>3 biological replicates. **(G)** Reproducibility of TimeLapse-seq libraries. Pearson correlation matrices of log-transformed gene-level read counts, shown as log1p (counts, across Timelapse-seq replicates of ΔDF1/ΔDF3, Tyr68His, Tyr153His, and ΔDF1-DF3+siDF2 cells, each paired with corresponding unlabeled control (ctrl) samples (62000 genes). Heatmap color indicates the Pearson correlation coefficient. **(H)** 4sU-dependent nucleotide conversion rate per replicate for ΔDF1/ΔDF3, Tyr68His, Tyr153His, and ΔDF1/ΔDF3+siDF2 cells. Upper bars indicate mutation rate at converted (peak, 4sU-labeled, p-new) positions; lower bars indicate background (unconverted, p-old) mutation rate. The clear separation between converted and background mutation rates across all replicates and conditions confirms successful and consistent 4sU incorporation and chemical conversion, supporting the k_deg estimates used in Figure 2. **(I)** Quantification of Δlog(k_deg) stratified by metagene region (5′UTR, n = 1,029; CDS, n = 4,123; 3′UTR, n = 5,029 transcripts) for Tyr68His, Tyr153His, and ΔDF1-DF3+siDF2 cells, corresponding to the metagene distribution shown in Figure 2E. Coding sequence (CDS)-localized transcripts showed significantly different Δk_deg compared with 3′UTR-localized transcripts in all three conditions (Tyr68His: p.adj = 1.1×10⁻⁴; Tyr153His: p.adj = 6.8×10⁻³; ΔDF1/ΔDF3+siDF2: p.adj = 5.8×10⁻¹⁸, Wilcoxon rank-sum test, BH-adjusted) *p < 0.05, **p < 0.01, ***p < 0.001, ****p < 0.0001.

**Figure S3:**
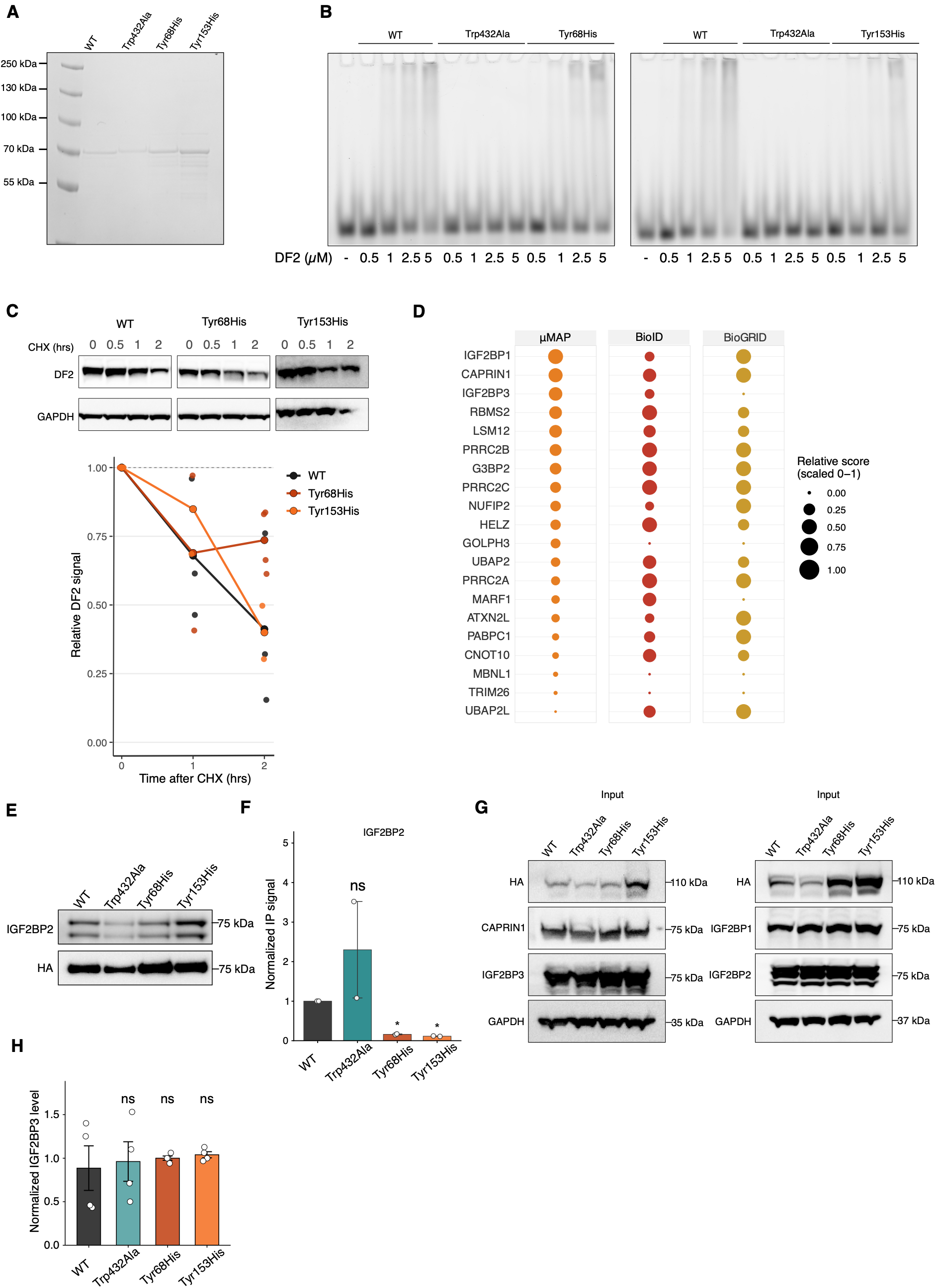
Biochemical and interaction validation of YTHDF2 IDR mutants. (**A**) Purification of YTHDF2 proteins. Gel code blue-stained SDS-PAGE gel showing recombinant YTHDF2 wild-type (WT), the m^6^A-binding-deficient YTHDF2-Trp432Ala, YTHDF2-Tyr68His, and YTHDF2-Tyr153His purified from E. coli. (**B**) Electrophoretic mobility shift assay (EMSA) measuring binding of recombinant YTHDF2 wild-type (WT), YTHDF2-Trp432Ala (m^6^A-binding-deficient control), and YTHDF2-Tyr68His (left) or YTHDF2-Tyr153His (right) to an m^6^A-containing RNA probe, across a titration of YTHDF2 protein (0, 0.5, 1, 2.5, 5 µM). **(C)** Cycloheximide (CHX) chase analysis of YTHDF2 protein stability. ΔDF1-3 cells stably expressing YTHDF2-WT, Tyr68His, or Tyr153His were treated with 100ug/mL CHX, and protein levels were analyzed at 0, 0.5, 1, and 2 hours. These time points were used because YTHDF2 has a short half-life ^45^. Top, representative immunoblot of YTHDF2 at 0, 0.5, 1, and 2 h after CHX treatment. GAPDH is used as a loading control because it has a half-life longer than 2 hours. Bottom, quantification of relative YTHDF2 signal (normalized to GAPDH and to time 0) over the CHX time course. Data are mean ± SEM from n = 2 biological replicates.(**D**) Cross-dataset comparison of the top endogenous YTHDF2 μMAP interactors against previously published YTHDF2 interactome datasets ^47,48,50^. Dot size indicates relative interaction score in each dataset. (**E**) BirA proximity labeling assay for IGF2BP2 association with YTHDF2 variants. Left, BioID proximity labeling in ΔDF1-3 cells transiently expressing BirA-tagged YTHDF2-HA-WT, YTHDF2-HA-Trp432Ala, YTHDF2-HA-Tyr68His, or YTHDF2-HA-Tyr153His. Streptavidin-enriched biotinylated proteins were immunoblotted for IGF2BP2 and HA. (**F**) Quantification of streptavidin pulldown signal for IGF2BP2, normalized to YTHDF2 construct expression and expressed relative to YTHDF2-HA-Trp432Ala. Data are mean ± SEM from n = 2 biological replicates. (**G**) Representative immunoblot showing the abundance of the indicated proteins in input lysates from cells expressing BirA-tagged YTHDF2-HA-WT, YTHDF2-HA-Trp432Ala, YTHDF2-HA-Tyr68His, or YTHDF2-HA-Tyr153His used for the pulldown experiments reported in Figure 3. (**H**) Quantification of IGF2BP3 protein levels in cells expressing BirA-tagged YTHDF2-HA-WT, YTHDF2-HA-Trp432Ala, YTHDF2-HA-Tyr68His, or YTHDF2-HA-Tyr153His, normalized to GAPDH. Data are mean ± SEM from n = 3-4 biological replicates as indicated.

**Figure S4:**
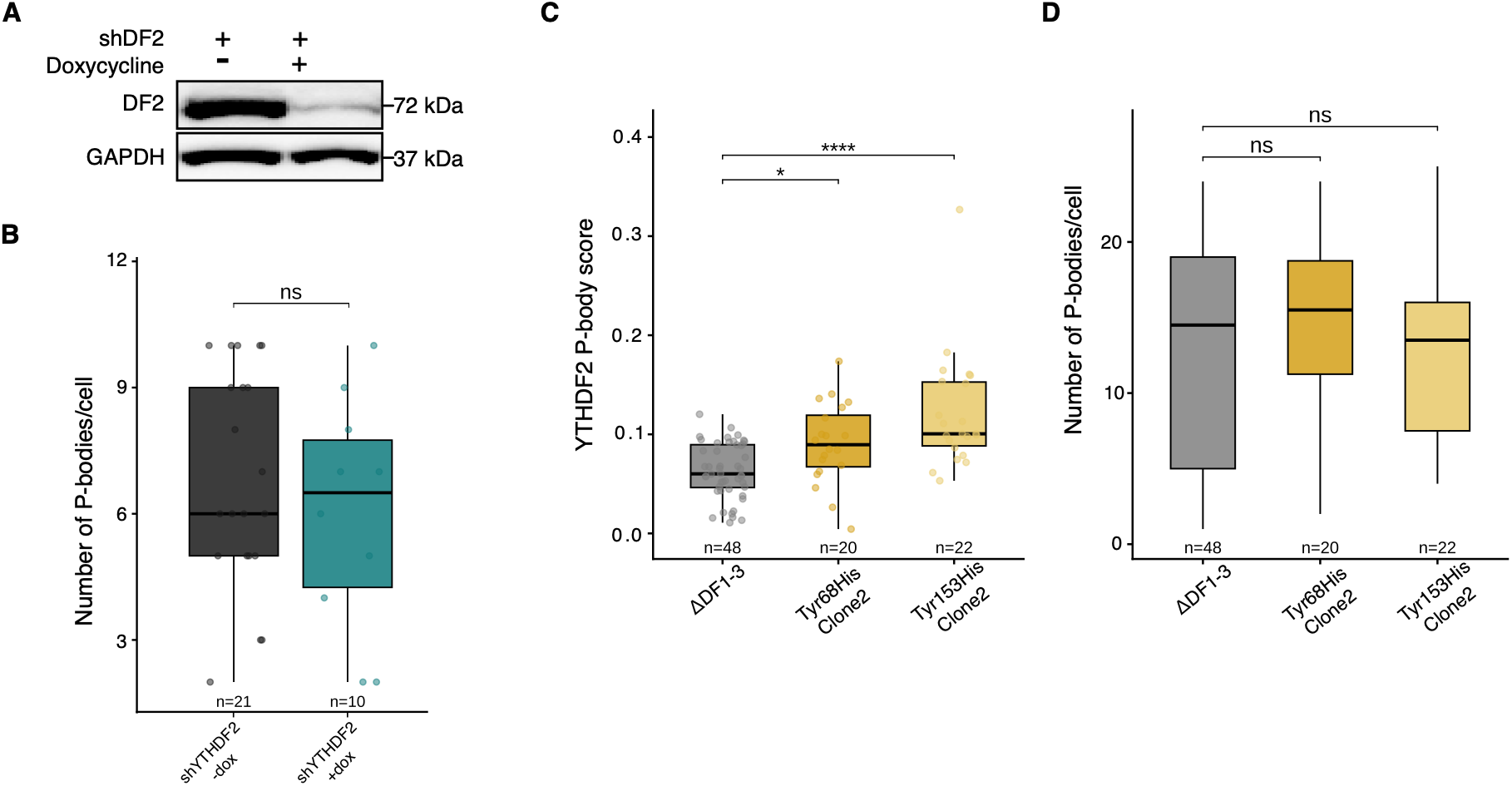
YTHDF2 depletion and additional analysis of YTHDF2 localization in wild-type and mutant lines. (**A**) Immunoblot of YTHDF2 and GAPDH in ΔDF1/ΔDF3 cells carrying a doxycycline-inducible shDF2 construct, cultured with or without 1ug/mL doxycycline, confirming efficient YTHDF2 depletion after 24 hours of treatment. (**B**) Quantification of P-body number per cell in ΔDF1/ΔDF3 cells carrying the doxycycline-inducible shDF2 construct, with or without doxycycline. YTHDF2 depletion did not significantly alter P-body number, indicating that P-body levels in this system are not driven by YTHDF2 abundance. Wilcoxon rank-sum test versus ΔDF1/ΔDF3, BH-adjusted. *p < 0.05, **p < 0.01, ***p < 0.001, ****p < 0.0001. (**C**) YTHDF2 enrichment at P-bodies in additional Tyr68His and Tyr153His clonal cell lines, extending the analysis shown in Figure 4B-C. Wilcoxon rank-sum test versus ΔDF1/ΔDF3, BH-adjusted. *p < 0.05, **p < 0.01, ***p < 0.001, ****p < 0.0001. (**D**) Quantification of P-body number per cell in the additional Tyr68His and Tyr153His clonal cell lines shown in (**C**). Wilcoxon rank-sum test versus ΔDF1/ΔDF3, BH-adjusted. *p < 0.05, **p < 0.01, ***p < 0.001, ****p < 0.0001.

## REFERENCES

1. Dominissini, D., Moshitch-Moshkovitz, S., Schwartz, S., Salmon-Divon, M., Ungar, L., Osenberg, S., Cesarkas, K., Jacob-Hirsch, J., Amariglio, N., Kupiec, M., et al. (2012). Topology of the human and mouse m6A RNA methylomes revealed by m6A-seq. Nature 485, 201–206. 10.1038/nature11112.

2. Liu, C., Sun, H., Yi, Y., Shen, W., Li, K., Xiao, Y., Li, F., Li, Y., Hou, Y., Lu, B., et al. (2023). Absolute quantification of single-base m6A methylation in the mammalian transcriptome using GLORI. Nat Biotechnol 41, 355–366. 10.1038/s41587-022-01487-9.

3. Meyer, K.D., Saletore, Y., Zumbo, P., Elemento, O., Mason, C.E., and Jaffrey, S.R. (2012). Comprehensive Analysis of mRNA Methylation Reveals Enrichment in 3′ UTRs and near Stop Codons. Cell 149, 1635–1646. 10.1016/j.cell.2012.05.003.

4. Roundtree, I.A., Evans, M.E., Pan, T., and He, C. (2017). Dynamic RNA Modifications in Gene Expression Regulation. Cell 169, 1187–1200. 10.1016/j.cell.2017.05.045.

5. Sendinc, E., and Shi, Y. (2023). RNA m6A methylation across the transcriptome. Molecular Cell 83, 428–441. 10.1016/j.molcel.2023.01.006.

6. Dierks, D., Shachar, R., Nir, R., Garcia-Campos, M.A., Uzonyi, A., Wiener, D., Toth, U., Rossmanith, W., Lasman, L., Slobodin, B., et al. (2025). Passive shaping of intra- and intercellular m6A dynamics via mRNA metabolism. eLife 13, RP100448. 10.7554/eLife.100448.

7. Sommer, S., Lavi, U., and Darnell, J.E. (1978). The absolute frequency of labeled N-6-methyladenosine in HeLa cell messenger RNA decreases with label time. J Mol Biol 124, 487–499. 10.1016/0022-2836(78)90183-3.

8. Du, H., Zhao, Y., He, J., Zhang, Y., Xi, H., Liu, M., Ma, J., and Wu, L. (2016). YTHDF2 destabilizes m6A-containing RNA through direct recruitment of the CCR4–NOT deadenylase complex. Nat Commun 7, 12626. 10.1038/ncomms12626.

9. Lasman, L., Krupalnik, V., Viukov, S., Mor, N., Aguilera-Castrejon, A., Schneir, D., Bayerl, J., Mizrahi, O., Peles, S., Tawil, S., et al. (2020). Context-dependent functional compensation between Ythdf m6A reader proteins. Genes Dev 34, 1373–1391. 10.1101/gad.340695.120.

10. Wang, X., Lu, Z., Gomez, A., Hon, G.C., Yue, Y., Han, D., Fu, Y., Parisien, M., Dai, Q., Jia, G., et al. (2014). N6-methyladenosine-dependent regulation of messenger RNA stability. Nature 505, 117–120. 10.1038/nature12730.

11. Zaccara, S., and Jaffrey, S.R. (2020). A Unified Model for the Function of YTHDF Proteins in Regulating m6A-Modified mRNA. Cell 181, 1582–1595.e18. 10.1016/j.cell.2020.05.012.

12. Zhu, T., Roundtree, I.A., Wang, P., Wang, X., Wang, L., Sun, C., Tian, Y., Li, J., He, C., and Xu, Y. (2014). Crystal structure of the YTH domain of YTHDF2 reveals mechanism for recognition of N6-methyladenosine. Cell Res 24, 1493–1496. 10.1038/cr.2014.152.

13. Cazzanelli, G., Dalle Vedove, A., Spagnolli, G., Terruzzi, L., Colasurdo, E., Boldrini, A., Patsilinakos, A., Sturlese, M., Grottesi, A., Biasini, E., et al. (2024). Pliability in the m6A-Binding Region Extends Druggability of YTH Domains. J Chem Inf Model 64, 1682–1690. 10.1021/acs.jcim.4c00051.

14. Li, F., Zhao, D., Wu, J., and Shi, Y. (2014). Structure of the YTH domain of human YTHDF2 in complex with an m6A mononucleotide reveals an aromatic cage for m6A recognition. Cell Res 24, 1490–1492. 10.1038/cr.2014.153.

15. Li, Y., Bedi, R.K., Moroz-Omori, E.V., and Caflisch, A. (2020). Structural and Dynamic Insights into Redundant Function of YTHDF Proteins. J Chem Inf Model 60, 5932–5935. 10.1021/acs.jcim.0c01029.

16. Wang, X., Lu, Z., Gomez, A., Hon, G.C., Yue, Y., Han, D., Fu, Y., Parisien, M., Dai, Q., Jia, G., et al. (2014). N6-methyladenosine-dependent regulation of messenger RNA stability. Nature 505, 117–120. 10.1038/nature12730.

17. Ries, R.J., Zaccara, S., Klein, P., Olarerin-George, A., Namkoong, S., Pickering, B.F., Patil, D.P., Kwak, H., Lee, J.H., and Jaffrey, S.R. (2019). m6A enhances the phase separation potential of mRNA. Nature 571, 424–428. 10.1038/s41586-019-1374-1.

18. Boo, S.H., Ha, H., Lee, Y., Shin, M.-K., Lee, S., and Kim, Y.K. (2022). UPF1 promotes rapid degradation of m6A-containing RNAs. Cell Rep 39, 110861. 10.1016/j.celrep.2022.110861.

19. Park, O.H., Ha, H., Lee, Y., Boo, S.H., Kwon, D.H., Song, H.K., and Kim, Y.K. (2019). Endoribonucleolytic Cleavage of m6A-Containing RNAs by RNase P/MRP Complex. Molecular Cell 74, 494–507.e8. 10.1016/j.molcel.2019.02.034.

20. Li, Q., Liu, J., Guo, L., Zhang, Y., Chen, Y., Liu, H., Cheng, H., Deng, L., Qiu, J., Zhang, K., et al. (2024). Decoding the interplay between m6A modification and stress granule stability by live-cell imaging. Sci Adv 10, eadp5689. 10.1126/sciadv.adp5689.

21. Zhang, F., Kang, Y., Wang, M., Li, Y., Xu, T., Yang, W., Song, H., Wu, H., Shu, Q., and Jin, P. (2018). Fragile X mental retardation protein modulates the stability of its m6A-marked messenger RNA targets. Hum Mol Genet 27, 3936–3950. 10.1093/hmg/ddy292.

22. Alberti, S., Gladfelter, A., and Mittag, T. (2019). Considerations and Challenges in Studying Liquid-Liquid Phase Separation and Biomolecular Condensates. Cell 176, 419–434. 10.1016/j.cell.2018.12.035.

23. Holehouse, A.S., and Alberti, S. (2025). Molecular determinants of condensate composition. Molecular Cell 85, 290–308. 10.1016/j.molcel.2024.12.021.

24. Cuella-Martin, R., Hayward, S.B., Fan, X., Chen, X., Huang, J.-W., Taglialatela, A., Leuzzi, G., Zhao, J., Rabadan, R., Lu, C., et al. (2021). Functional interrogation of DNA damage response variants with base editing screens. Cell 184, 1081–1097.e19. 10.1016/j.cell.2021.01.041.

25. Hanna, R.E., Hegde, M., Fagre, C.R., DeWeirdt, P.C., Sangree, A.K., Szegletes, Z., Griffith, A., Feeley, M.N., Sanson, K.R., Baidi, Y., et al. (2021). Massively parallel assessment of human variants with base editor screens. Cell 184, 1064–1080.e20. 10.1016/j.cell.2021.01.012.

26. Lue, N.Z., and Liau, B.B. (2023). Base editor screens for in situ mutational scanning at scale. Mol Cell 83, 2167–2187. 10.1016/j.molcel.2023.06.009.

27. Gaudelli, N.M., Komor, A.C., Rees, H.A., Packer, M.S., Badran, A.H., Bryson, D.I., and Liu, D.R. (2017). Programmable base editing of A•T to G•C in genomic DNA without DNA cleavage. Nature 551, 464–471. 10.1038/nature24644.

28. Komor, A.C., Kim, Y.B., Packer, M.S., Zuris, J.A., and Liu, D.R. (2016). Programmable editing of a target base in genomic DNA without double-stranded DNA cleavage. Nature 533, 420–424. 10.1038/nature17946.

29. Nishimasu, H., Shi, X., Ishiguro, S., Gao, L., Hirano, S., Okazaki, S., Noda, T., Abudayyeh, O.O., Gootenberg, J.S., Mori, H., et al. (2018). Engineered CRISPR-Cas9 nuclease with expanded targeting space. Science 361, 1259–1262. 10.1126/science.aas9129.

30. Sangree, A.K., Griffith, A.L., Szegletes, Z.M., Roy, P., DeWeirdt, P.C., Hegde, M., McGee, A.V., Hanna, R.E., and Doench, J.G. (2022). Benchmarking of SpCas9 variants enables deeper base editor screens of BRCA1 and BCL2. Nat Commun 13, 1318. 10.1038/s41467-022-28884-7.

31. Rodrigues, C.H.M., Pires, D.E.V., and Ascher, D.B. (2021). DynaMut2: Assessing changes in stability and flexibility upon single and multiple point missense mutations. Protein Science 30, 60–69. 10.1002/pro.3942.

32. Pires, D.E.V., and Ascher, D.B. (2017). mCSM–NA: predicting the effects of mutations on protein–nucleic acids interactions. Nucleic Acids Res 45, W241–W246. 10.1093/nar/gkx236.

33. Lancaster, A.K., Nutter-Upham, A., Lindquist, S., and King, O.D. (2014). PLAAC: a web and command-line application to identify proteins with prion-like amino acid composition. Bioinformatics 30, 2501–2502. 10.1093/bioinformatics/btu310.

34. Holehouse, A.S., Das, R.K., Ahad, J.N., Richardson, M.O.G., and Pappu, R.V. (2017). CIDER: Resources to Analyze Sequence-Ensemble Relationships of Intrinsically Disordered Proteins. Biophys J 112, 16–21. 10.1016/j.bpj.2016.11.3200.

35. Mészáros, B., Erdos, G., and Dosztányi, Z. (2018). IUPred2A: context-dependent prediction of protein disorder as a function of redox state and protein binding. Nucleic Acids Res 46, W329–W337. 10.1093/nar/gky384.

36. Vendruscolo, M., and Fuxreiter, M. (2026). FuzDrop: sequence-based prediction of the propensity of proteins for liquid–liquid phase separation and aggregation. Nat Protoc 21, 2016–2042. 10.1038/s41596-025-01267-0.

37. Hornbeck, P.V., Kornhauser, J.M., Tkachev, S., Zhang, B., Skrzypek, E., Murray, B., Latham, V., and Sullivan, M. (2012). PhosphoSitePlus: a comprehensive resource for investigating the structure and function of experimentally determined post-translational modifications in man and mouse. Nucleic Acids Res 40, D261–270. 10.1093/nar/gkr1122.

38. Chen, Y., Wan, R., Zou, Z., Lao, L., Shao, G., Zheng, Y., Tang, L., Yuan, Y., Ge, Y., He, C., et al. (2023). O-GlcNAcylation determines the translational regulation and phase separation of YTHDF proteins. Nat Cell Biol 25, 1676–1690. 10.1038/s41556-023-01258-x.

39. Zou, Z., Sepich-Poore, C., Zhou, X., Wei, J., and He, C. (2023). The mechanism underlying redundant functions of the YTHDF proteins. Genome Biol 24, 17. 10.1186/s13059-023-02862-8.

40. Vock, I.W., Mabin, J.W., Machyna, M., Zhang, A., Hogg, J.R., and Simon, M.D. (2025). Expanding and improving analyses of nucleotide recoding RNA-seq experiments with the EZbakR suite. PLOS Computational Biology 21, e1013179. 10.1371/journal.pcbi.1013179.

41. Murakami, S., Olarerin-George, A.O., Liu, J.F., Zaccara, S., Hawley, B., and Jaffrey, S.R. (2025). m6A alters ribosome dynamics to initiate mRNA degradation. Cell 188, 3728–3743.e20. 10.1016/j.cell.2025.04.020.

42. Zhou, Y., Ćorović, M., Hoch-Kraft, P., Meiser, N., Mesitov, M., Körtel, N., Back, H., Naarmann-de Vries, I.S., Katti, K., Obrdlík, A., et al. (2024). m6A sites in the coding region trigger translation-dependent mRNA decay. Mol Cell 84, 4576–4593.e12. 10.1016/j.molcel.2024.10.033.

43. Capraro, F., Abis, G., Incocciati, A., Simpson, P.J., Karimzadeh, M., Masino, L., Barley, A., Bui, T.T.T., Kelly, G., Goodarzi, H., et al. (2026). An intrinsically disordered region mediates RNA-binding selectivity and cellular activities of LARP6. Nat Commun 17, 2939. 10.1038/s41467-026-69789-z.

44. Qiu, C., Zhang, Z., Wine, R.N., Campbell, Z.T., Zhang, J., and Hall, T.M.T. (2023). Intra- and inter-molecular regulation by intrinsically-disordered regions governs PUF protein RNA binding. Nat Commun 14, 7323. 10.1038/s41467-023-43098-1.

45. Li, J., Zhou, W., Zhang, J., Ma, L., Lv, Z., Geng, Y., Chen, X., and Li, J. (2025). O-GlcNAcylation of YTHDF2 antagonizes ERK-dependent phosphorylation and inhibits lung carcinoma. Fundamental Research 5, 2388–2396. 10.1016/j.fmre.2024.07.003.

46. Davey, N.E., Van Roey, K., Weatheritt, R.J., Toedt, G., Uyar, B., Altenberg, B., Budd, A., Diella, F., Dinkel, H., and Gibson, T.J. (2012). Attributes of short linear motifs. Mol. BioSyst. 8, 268–281. 10.1039/c1mb05231d.

47. Go, C.D., Knight, J.D.R., Rajasekharan, A., Rathod, B., Hesketh, G.G., Abe, K.T., Youn, J.-Y., Samavarchi-Tehrani, P., Zhang, H., Zhu, L.Y., et al. (2021). A proximity-dependent biotinylation map of a human cell. Nature 595, 120–124. 10.1038/s41586-021-03592-2.

48. Geri, J.B., Oakley, J.V., Reyes-Robles, T., Wang, T., McCarver, S.J., White, C.H., Rodriguez-Rivera, F.P., Parker, D.L., Hett, E.C., Fadeyi, O.O., et al. (2020). Microenvironment mapping via Dexter energy transfer on immune cells. Science 367, 1091–1097. 10.1126/science.aay4106.

49. Stadler, C., Skogs, M., Brismar, H., Uhlén, M., and Lundberg, E. (2010). A single fixation protocol for proteome-wide immunofluorescence localization studies. Journal of Proteomics 73, 1067–1078. 10.1016/j.jprot.2009.10.012.

50. Oughtred, R., Rust, J., Chang, C., Breitkreutz, B.-J., Stark, C., Willems, A., Boucher, L., Leung, G., Kolas, N., Zhang, F., et al. (2021). The BioGRID database: A comprehensive biomedical resource of curated protein, genetic, and chemical interactions. Protein Science 30, 187–200. 10.1002/pro.3978.

51. Youn, J.-Y., Dunham, W.H., Hong, S.J., Knight, J.D.R., Bashkurov, M., Chen, G.I., Bagci, H., Rathod, B., MacLeod, G., Eng, S.W.M., et al. (2018). High-Density Proximity Mapping Reveals the Subcellular Organization of mRNA-Associated Granules and Bodies. Mol Cell 69, 517–532.e11. 10.1016/j.molcel.2017.12.020.

52. Chen, B., Huang, R., Xia, T., Wang, C., Xiao, X., Lu, S., Chen, X., Ouyang, Y., Deng, X., Miao, J., et al. (2023). The m6A reader IGF2BP3 preserves NOTCH3 mRNA stability to sustain Notch3 signaling and promote tumor metastasis in nasopharyngeal carcinoma. Oncogene 42, 3564–3574. 10.1038/s41388-023-02865-6.

53. Huang, H., Weng, H., Sun, W., Qin, X., Shi, H., Wu, H., Zhao, B.S., Mesquita, A., Liu, C., Yuan, C.L., et al. (2018). Recognition of RNA N6-methyladenosine by IGF2BP Proteins Enhances mRNA Stability and Translation. Nat Cell Biol 20, 285–295. 10.1038/s41556-018-0045-z.

54. Branon, T.C., Bosch, J.A., Sanchez, A.D., Udeshi, N.D., Svinkina, T., Carr, S.A., Feldman, J.L., Perrimon, N., and Ting, A.Y. (2018). Efficient proximity labeling in living cells and organisms with TurboID. Nat Biotechnol 36, 880–887. 10.1038/nbt.4201.

55. Huang, H., Weng, H., Sun, W., Qin, X., Shi, H., Wu, H., Zhao, B.S., Mesquita, A., Liu, C., Yuan, C.L., et al. (2018). Recognition of RNA N6-methyladenosine by IGF2BP proteins enhances mRNA stability and translation. Nat Cell Biol 20, 285–295. 10.1038/s41556-018-0045-z.

56. Fu, Y., and Zhuang, X. (2020). m6A-binding YTHDF proteins promote stress granule formation. Nat Chem Biol 16, 955–963. 10.1038/s41589-020-0524-y.

57. Yu, J.H., Yang, W.-H., Gulick, T., Bloch, K.D., and Bloch, D.B. (2005). Ge-1 is a central component of the mammalian cytoplasmic mRNA processing body. RNA 11, 1795– 1802. 10.1261/rna.2142405.

58. Khong, A., Matheny, T., Jain, S., Mitchell, S.F., Wheeler, J.R., and Parker, R. (2017). The Stress Granule Transcriptome Reveals Principles of mRNA Accumulation in Stress Granules. Molecular Cell 68, 808–820.e5. 10.1016/j.molcel.2017.10.015.

59. Kedersha, N.L., Gupta, M., Li, W., Miller, I., and Anderson, P. (1999). RNA-Binding Proteins Tia-1 and Tiar Link the Phosphorylation of Eif-2α to the Assembly of Mammalian Stress Granules. J Cell Biol 147, 1431–1442. 10.1083/jcb.147.7.1431.

60. Duan, M., Liu, H., Xu, S., Yang, Z., Zhang, F., Wang, G., Wang, Y., Zhao, S., and Jiang, X. (2024). IGF2BPs as novel m6A readers: Diverse roles in regulating cancer cell biological functions, hypoxia adaptation, metabolism, and immunosuppressive tumor microenvironment. Genes & Diseases 11, 890–920. 10.1016/j.gendis.2023.06.017.

61. Ramesh-Kumar, D., and Guil, S. (2022). The IGF2BP family of RNA binding proteins links epitranscriptomics to cancer. Semin Cancer Biol 86, 18–31. 10.1016/j.semcancer.2022.05.009.

62. Zhou, X., Li, G., Su, X., Wang, Y., Liu, L., Yang, C., Xu, Y., He, J., and Zhang, J. (2026). The role of m6A reader IGF2BPs in tumor metabolic reprogramming. Biochem Pharmacol 243, 117452. 10.1016/j.bcp.2025.117452.

63. Parker, R., and Sheth, U. (2007). P Bodies and the Control of mRNA Translation and Degradation. Molecular Cell 25, 635–646. 10.1016/j.molcel.2007.02.011.

64. Standart, N., and Weil, D. (2018). P-Bodies: Cytosolic Droplets for Coordinated mRNA Storage. Trends in Genetics 34, 612–626. 10.1016/j.tig.2018.05.005.

65. Chan, W.Y., Jin, E., Cross, H., Masino, L., Ibrahim, F., Albihlal, W.S., Berger, D., Boot, J., Michrowska, A., Mouilleron, S., et al. (2026). Cellular context restricts a promiscuous m6A reader IDR to a single functional effector interface for mRNA decay. Preprint at bioRxiv, 10.64898/2026.08.04.742832 https://doi.org/10.64898/2026.08.04.742832.

66. Hofweber, M., and Dormann, D. (2019). Friend or foe-Post-translational modifications as regulators of phase separation and RNP granule dynamics. J Biol Chem 294, 7137–7150. 10.1074/jbc.TM118.001189.

67. Zou, Z., Wei, J., Chen, Y., Kang, Y., Shi, H., Yang, F., Shi, Z., Chen, S., Zhou, Y., Sepich-Poore, C., et al. (2023). FMRP phosphorylation modulates neuronal translation through YTHDF1. Molecular Cell 83, 4304–4317.e8. 10.1016/j.molcel.2023.10.028.

68. Wang, X., Codreanu, S.G., Wen, B., Li, K., Chambers, M.C., Liebler, D.C., and Zhang, B. (2018). Detection of Proteome Diversity Resulted from Alternative Splicing is Limited by Trypsin Cleavage Specificity*. Molecular C Cellular Proteomics 17, 422–430. 10.1074/mcp.RA117.000155.

69. Hellman, L.M., and Fried, M.G. (2007). Electrophoretic mobility shift assay (EMSA) for detecting protein–nucleic acid interactions. Nat Protoc 2, 1849–1861. 10.1038/nprot.2007.249.

70. Schofield, J.A., Duffy, E.E., Kiefer, L., Sullivan, M.C., and Simon, M.D. (2018). TimeLapse-seq: adding a temporal dimension to RNA sequencing through nucleoside recoding. Nat Methods 15, 221–225. 10.1038/nmeth.4582.

